# The photoreceptor UVR8 mediates the perception of both UV-B and UV-A wavelengths up to 350 nm of sunlight with responsivity moderated by cryptochromes

**DOI:** 10.1101/2020.01.21.913814

**Authors:** Neha Rai, Andrew O’Hara, Daniel Farkas, Omid Safronov, Khuanpiroon Ratanasopa, Fang Wang, Anders V. Lindfors, Gareth I. Jenkins, Tarja Lehto, Jarkko Salojärvi, Mikael Brosché, Åke Strid, Pedro J. Aphalo, Luis O. Morales

## Abstract

The photoreceptors UV RESISTANCE LOCUS 8 (UVR8) and CRYPTOCHROMES 1 and 2 (CRYs) play major roles in the perception of UV-B (280–315 nm) and UV-A/blue radiation (315–500 nm), respectively. However, it is poorly understood how they function in sunlight. The roles of UVR8 and CRYs were assessed in a factorial experiment with *Arabidopsis thaliana* wild-type and photoreceptor mutants exposed to sunlight for 6 h or 12 h under five types of filters with cut-offs in UV and blue-light regions. Transcriptome-wide responses triggered by UV-B and UV-A wavelengths shorter than 350 nm (UV-A_sw_) required UVR8 whereas those induced by blue and UV-A wavelengths longer than 350 nm (UV-A_lw_) required CRYs. UVR8 modulated gene expression in response to blue light while lack of CRYs drastically enhanced gene expression in response to UV-B and UV-A_sw_. These results agree with our estimates of photons absorbed by these photoreceptors in sunlight and with *in vitro* monomerization of UVR8 by wavelengths up to 335 nm. Motif enrichment analysis predicted complex signaling downstream of UVR8 and CRYs. Our results highlight that it is important to use UV waveband definitions specific to plants’ photomorphogenesis as is routinely done in the visible region.

## 1 INTRODUCTION

Sunlight regulates plant growth, development and acclimation to the environment, while responses to specific wavelengths are regulated by different photoreceptors. The contribution of different wavelengths of sunlight to plant responses depends both on the optical properties of the photoreceptors and on the spectrum and photon irradiance of the incident radiation. Research under controlled conditions has shown that the photoreceptors UV RESISTANCE LOCUS 8 (UVR8) and CRYPTOCHROMES 1 and 2 (CRYs) play major roles in the perception of UV-B (ground level 290–315 nm) and UV-A/blue radiation (315–500 nm), respectively (Ahmad & Cashmore, 1993; Lin, 2000; Yu *et al*., 2010; Rizzini *et al*., 2011). However, in sunlight, the irradiances of photosynthetically active radiation (PAR, 400–700 nm) and UV-A (315–400 nm) relative to UV-B are much higher than those normally used in controlled environments making it necessary to assess which wavelengths are effectively perceived by UVR8 and CRYs in sunlight.

Indoor and outdoor experiments with UV-B irradiances similar to those in sunlight have shown that UVR8 can regulate transcript abundance of hundreds of genes, including those involved in UV protection, photo-repair of UV-B-induced DNA damage, oxidative stress and several transcription factors (TFs) shared with other signaling pathways (Brown *et al*., 2005; Favory *et al*., 2009; Morales *et al*., 2013). The first step in this response is the absorption of UV-B photons by UVR8 which triggers a change in conformation from homodimer to monomer enhancing its accumulation in the nucleus (Brown *et al*., 2005; Kaiserli & Jenkins, 2007; Rizzini *et al*., 2011). Downstream signaling depends on UVR8 monomers binding to CONSTITUTIVE PHOTOMORPHOGENIC 1 (COP1) (Favory *et al*., 2009), thereby inactivating the E3 ubiquitin ligase activity of COP1. The UVR8-COP1 association stabilizes the TF ELONGATED HYPOCOTYL 5 (HY5), a master regulator of photomorphogenesis in plants (Favory *et al*., 2009; Huang *et al*., 2013; Gangappa & Botto, 2016). Both HY5 and HY5 HOMOLOG (HYH) have been identified as key TFs regulating the expression of most genes responding to UV-B through UVR8 signaling (Brown & Jenkins, 2008; Favory *et al*., 2009). Moreover, UVR8 directly interacts with WRKY DNA-BINDING PROTEIN 36 (WRKY36), BRI1-EMS-SUPPRESSOR1 (BES1) and BES1-INTERACTING MYC-

LIKE 1 (BIM1) TFs to regulate transcription (Liang *et al*., 2018; Yang *et al*., 2018). UV-B radiation induces the expression of genes encoding REPRESSOR OF UV-B PHOTOMORPHOGENESIS 1 (RUP1) and RUP2 proteins, which interact with UVR8 directly and convert the active monomers into homodimers, thereby decreasing the abundance of UVR8 monomers through negative feedback (Gruber *et al*., 2010; Heijde & Ulm, 2013). Because indoor studies are often done using unrealistically high UV-B:PAR and low UV-A:PAR ratios, the participation of UVR8 in the regulation of the transcriptome in response to UV-B, UV-A and blue wavelengths of solar radiation remains uncertain.

Our understanding of the molecular mechanisms underpinning CRY-mediated responses to blue light is far better than for those to UV-A. In indoor experiments with blue light, CRYs have been found to regulate photomorphogenesis and expression of genes involved in light signaling, photosynthetic light reaction, the Calvin cycle, phenylpropanoid metabolic pathway and stress response (Ohgishi *et al*., 2004; Kleine *et al*., 2007). Upon absorption of blue light photons, CRYs alter conformation from monomers to homodimers and oligomers (Wang *et al*., 2016). CRYs homodimers or oligomers interact with COP1 and SUPPRESSOR OF PHY A (SPA) proteins through various mechanisms (Yang *et al*., 2001; Wang *et al*., 2001; Lian *et al*., 2011; Liu *et al*., 2011b; Zuo *et al*., 2011; Podolec & Ulm, 2018). These interactions stabilize HY5 abundance and consequent transcriptional regulation (Liu *et al*., 2011a; Yang *et al*., 2017; Podolec & Ulm, 2018). CRYs also interact directly with TFs such as BES1, BIM1, CRYPTOCHROME-INTERACTING basic helix-loop-helix 1 (CIB1), PHYTOCHROME-INTERACTING FACTOR 4 (PIF4) and PIF5 regulating transcription (Liu *et al*., 2008; Pedmale *et al*., 2016; Wang *et al*., 2018). Thus, both UVR8 and CRY signaling share some TFs such as HY5, HYH and BES1 suggesting the possibility of crosstalk. Despite these studies under controlled conditions, the participation of CRYs in the regulation of the transcriptome in sunlight remains poorly understood due to the unrealistic light conditions used.

As photoreceptors have broad peaks of absorption, their absorption spectra partially overlap. Both CRYs and UVR8 absorb in the UV-B region, while CRYs also absorb strongly at longer wavelengths (Banerjee *et al*., 2007; Christie *et al*., 2012; Yang *et al*., 2015). Given the overlapping spectra of photoreceptors and the fact that plants are simultaneously exposed to all wavelengths of sunlight, only research in sunlight can assess the roles of photoreceptors in nature. The available absorption spectrum for the UVR8 molecule covers the UV-C and UV-B regions, extending only 15 nm into the UV-A (Christie *et al*., 2012) making even speculations about the possible role of UVR8 in the UV-A region uncertain. Furthermore, our previous studies indicated that different regions within UV-A could trigger different responses to metabolite accumulation and transcript abundance of selected genes (Siipola *et al*., 2015; Rai *et al*., 2019). Besides, results from an indoor experiment suggested the participation of UVR8 in flavonoid accumulation in response to UV-A from LEDs (Brelsford *et al*., 2018). However, it is not yet clear which UV-A wavelengths are perceived through UVR8 and which ones through CRYs.

To assess the roles of UVR8 and CRYs in the perception of solar UV-B, UV-A and blue radiation, we measured transcriptome-wide responses in Arabidopsis plants exposed to sunlight. We also measured the *in vitro* absorption spectrum of UVR8 in the UV-C, UV-B, UV-A and visible regions (250–500 nm), the *in vitro* monomerization of UVR8 by different UV-A wavelengths, and estimated numbers of sunlight photons absorbed by UVR8 and CRYs. We tested four hypotheses: 1) the perception of solar UV-B and UV-A wavelengths up to 350 nm (UV-A_sw_) is through UVR8, 2) the perception of solar UV-A wavelengths above 350 nm (UV-A_lw_) and blue light is through CRYs, 3) crosstalk between UV-B/UV-A_sw_ and UV-A_lw_/blue light signaling is asymmetric in plants exposed to sunlight, and 4) gene expression responses to different wavelengths of sunlight are coordinated by multiple TFs resulting in multiple patterns of expression.

## 2 MATERIALS AND METHODS

### 2.1 Plant material and treatments

*Arabidopsis thaliana* ecotype Landsberg *erecta* (L*er*) and the photoreceptor mutants *uvr8-2* (Brown *et al*., 2005) and *cry1cry2* (Neff & Chory, 1998) were used. The *uvr8-2* genotype carries a mutation in the C-terminus of the UVR8 protein that impairs signaling in response to UV-B (Cloix *et al*., 2012). The *cry1cry2* genotype carries null mutations for CRY1 and CRY2 protein and consequently is impaired in blue light perception through CRYs (Mazzella *et al*., 2001). Seeds from different genotypes used in the experiments were previously grown and harvested under the same growth conditions. Seeds were sown in plastic pots (8 cm × 8 cm) containing a 1:1 mixture of peat and vermiculite and kept in darkness at 4 °C for 3 d. Subsequently, the pots were transferred to controlled-environment growth room at 23 °C:19 °C and 70%:90% relative humidity (light:dark) under 12 h photoperiod with 280 µmol m^−2^ s^−1^ white light irradiance, 12 mol day^−1^ (Osram T8 L 36W/865 Lumilux). Rosco filter E-color 226 was used to block the small amount of UV-A radiation emitted by the lamps. Four seedlings of the same genotype were transplanted into each plastic pot (8 cm × 8 cm). After transplanting, plants were kept for 14 d in the same growth room and conditions.

For exposure to sunlight, plants were moved to the field (Viikki campus, University of Helsinki, 60°13′N, 25°1′E) on 21 August 2014 between 07:30 and 08:15. Five treatments were created with different plastic sheets (3 mm thick) and a film (0.12 mm thick) used as long-pass optical filters to selectively exclude different wavebands of the UV and blue regions (Figure 1, Methods S1). The filters were kept 10–15 cm above the top of the plants, on their south and north edges, respectively. Their transmittance was measured with a spectrophotometer (model 8453, Agilent, Waldbronn, Germany, Figure 1). Treatments were randomly assigned within four blocks (biological replicates). One tray was kept under one filter and there was one filter of each type per block. Each tray contained two pots per genotype, positioned at random within the trays, for sampling after 6 h and 12 h.

**Figure 1.**
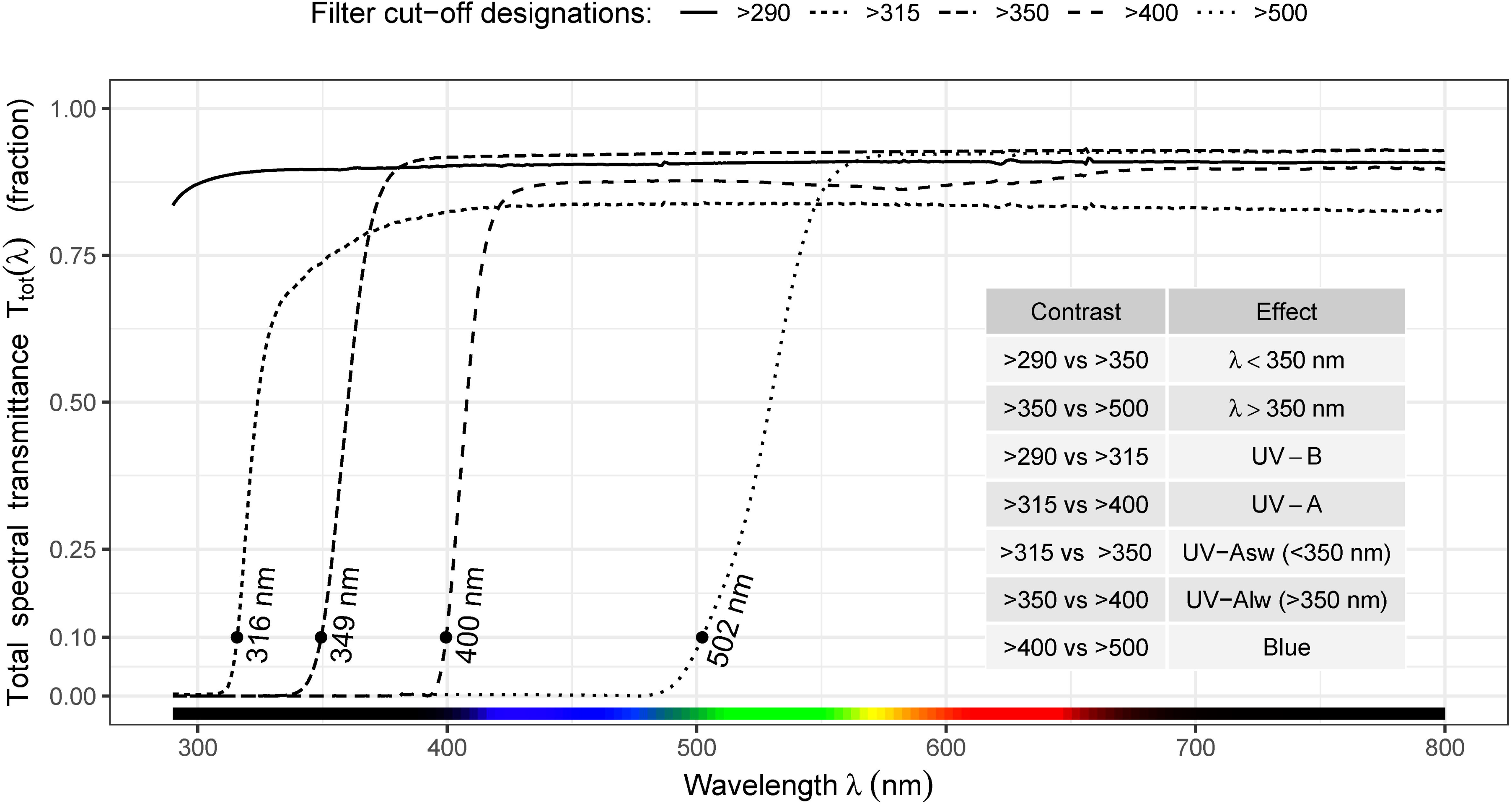
Transmittance of filters used in the outdoor experiment and the statistical contrasts between pairs of filter treatments used to assess the effects of different ranges of wavelengths in solar radiation.

### 2.2 Light conditions and sampling outdoors

Hourly solar spectra at ground level were modeled for the 12 h of exposure period using a radiation transfer model (libradtran, Emde *et al*., 2016) and cloudiness estimates derived from global radiation measurements (Lindfors *et al*., 2009). Figure S1 shows the solar spectrum at different times of the day when plants were moved outdoors. Figure S2 shows the hourly mean photon irradiance of UV-B (290–315 nm), UV-A_sw_ (315–350 nm), UV-A long-wavelength (UV-A_lw_ 350– 400 nm), blue (400–500 nm) and PAR (400–700 nm), for the broader wavebands λ < 350 nm (290– 350 nm) and λ > 350 nm (350–500 nm). The hourly solar spectra were convoluted by the spectral transmittance of each filter to estimate the spectrum the plants were exposed to. Then these spectra were convoluted by the *in vitro* spectral absorptance of UVR8 (see Results) or of light-adapted CRY2 (Banerjee *et al*., 2007) to estimate the relative numbers of photons absorbed by UVR8 and CRY2 in each treatment. Plotting and calculations on the simulated spectra were done in R (R Core Team, 2018, Aphalo, 2015).

Samples were collected after 6 h and 12 h of exposure to sunlight (13:30–14:15, 19:30–20:20) by block with treatments and genotypes in random order within each block. Each biological sample consisted of leaves from four pooled rosettes from the same pot, which were immediately frozen in liquid nitrogen and later stored at –80 °C. Each pooled sample was ground with mortar and pestle in liquid nitrogen.

### 2.3 RNA sequencing

Total RNA was extracted from ground leaf samples with a GeneJET Plant RNA Purification Kit following manufacturer’s guidelines (Thermo Fisher Scientific, Vilnius, Lithuania). RNA quality was checked with Agilent 2100 Bioanalyzer (Santa Clara, CA, USA) and RNA concentration was measured with ND-1000 Spectrophotometer (NanoDrop Technologies, Thermo Fisher Scientific, Waltham, MA, USA). For RNA-seq measurements, RNA extracts from two pairs of biological replicates were combined into two pooled replicates. Libraries were constructed using TruSeq Stranded mRNA Sample PrepKit (Illumina, San Diego, CA, USA) following manufacturer’s instructions. The library concentration was measured using Qubit Fluorometer (Life Technologies, Carlsbad, CA, USA), and quality and size were checked by Fragment Analyzer (Agilent Technologies). Libraries were sequenced on NextSeq 500 (Illumina) generating single end 75 bp reads. RNA-seq raw data was deposited at Gene Expression Omnibus (accession number GSE117199).

RNA-seq data analysis was done using the JAVA-based client-server system, Chipster (Kallio *et al*., 2011) and in R. The quality of raw reads was checked with FastQC (Andrews, 2014). Removal of adapter sequences, trimming and cropping of the reads were done using Trimmomatic-0.33 (Bolger *et al*., 2014) in single-end mode. The bases with a Phred score < 20 were trimmed from the ends of the reads, and the reads shorter than 30 bases were removed from the analysis (-Phred33, TRAILING:20, MINLEN:30).

Filtered reads were mapped to the Arabidopsis transcript reference database AtRTD2 (Zhang *et al*., 2017) using Kallisto V-0.43.0 (CMD:quant) (Bray *et al*., 2016) with 4000 bootstrap sets. The raw count tables for the two pooled replicates were obtained as the mean of the bootstrap runs. Genes with less than five counts in all 21 filter-treatment × genotype combinations were removed. The count tables were analyzed for differential gene expression with edgeR 3.24.3 (Robinson *et al*., 2010). The glmLRT (McCarthy *et al*., 2012) method was used to fit the statistical model separately to data from each genotype. Differentially expressed genes (DEGs) across treatments and photoreceptor mutants under selected pairwise contrasts were assessed with method decideTestsDGE using Benjamini-Hochberg FDR correction of *P*-values, with FDR ≤ 0.05. In a separate step, |logFC| > log_2_(1.5) was used as threshold. Effects of wavebands were assessed by comparing responses between pairs of filters as described above (Figure 1).

Function plotMDS from package limma 3.38.3 (Ritchie *et al*., 2015) was used to carry out dimensionality reduction to test for consistency between replicates. To compare RNA-seq and qRT-PCR estimates of transcript abundance, estimates from RNA-seq were re-expressed relative to L*er* UV0 to match qRT-PCR data and major axis regression applied (R package lmodel2 1.7-3, Legendre, 2018).

Enrichment of Kyoto Encyclopedia of Genes and Genomes (KEGG) pathways was assessed using function topKEGG (edgeR 3.24.3) using pathway definitions downloaded from http://rest.kegg.jp on 10 April 2019. Pathways whose definition included at least 15 but not more than 250 genes were included in the analysis. The enrichments were tested for all lists of DEGs from the contrast tests described above, and all the genes expressed in our experiment were used as background. Conditions used to assess significance of pathway enrichment were *P-*value cut-off of 0.01 and at least 1/3 of pathway genes differentially expressed. All pathways fulfilling both conditions in at least one treatment contrast are reported.

### 2.4 Cis-motif enrichment

Transcription factor binding motifs were collected in the form of position-specific weight matrices (PSWMs) from JASPAR 2018 (Khan *et al*., 2018) and Cistrome (O’Malley *et al*., 2016) databases. In addition to PSWMs we analyzed a previously collected set of binding motifs (Blomster *et al*., 2011). For each contrast within each genotype, the enrichment analyses were run on the lists of upregulated and downregulated genes from RNA-seq analysis. The promoter sequences 1000 base pairs upstream of transcription start sites of all Arabidopsis genes were scanned for binding motifs. The regular expressions were matched in R, and PSWM hits were identified using MEME (Ambrosini *et al*., 2018) with default thresholds. The significance of the overlap between the gene lists and motif occurrences was then tested with Fisher exact test, followed by FDR correction using Benjamini-Hochberg correction. Based on the motifs enriched in each of the separate lists of up or downregulated genes for the 12 waveband × genotype combinations, we identified putative TFs which could regulate the expression of those genes. Clustering was done over the obtained lists of TFs, based on the adjusted *P*-values for enrichment in each contrast and genotype combination. For visualization, the *P*-values were first restricted to the range of 1 to 10^−4^ by converting all smaller values to 10^−4^, and subsequently applying a log_10_ transformation. Clustering and plotting of the heatmaps was done with R package pheatmap 1.0.12 (Kolde, 2019). The cut point at 12 clusters was subjectively chosen for plotting; cutting introduces visual breaks without changing the cluster tree or the heatmap in any other way. The ordering of the tree was done with R package dendsort 0.3.3 (Sakai *et al*., 2014).

### 2.5 Quantitative real-time PCR and data analysis

Transcript abundance of selected genes was measured with qRT-PCR from four biological replicates and collected both at 6 h and 12 h. At 6 h the same RNA extracts were used for qRT-PCR as for RNA-seq, but without pooling. The qRT-PCR was done according to Rai *et al*. (2019) using primers listed in Table S1. In every run, normalized expression values were scaled to sample L*er* λ > 400 nm, log_10_ transformed, and exported from qbase^PLUS^ for statistical analyses in R. Linear mixed-effect models with block as a random factor were fitted using function lme from package ‘nlme’ 3.1-137 (Pinheiro *et al*., 2018). Factorial ANOVA was used to assess the significance of the main effects (treatment, genotype and exposure time) and of the interactions (treatment × genotype, treatment × exposure time, genotype × exposure time, and treatment × genotype × exposure time). Function fit.contrast from package gmodels 2.18.1 (Warnes *et al*., 2018) was used to fit pairwise contrasts defined *a priori* and *P*-values adjusted with function p.adjust in R (Holm, 1979). Figures were plotted using R package ggplot2 3.1.0 (Wickham, 2009).

### 2.6 In vitro absorption spectra of UVR8 protein

Recombinant UVR8 was produced and isolated from *Escherichia coli* with small variations from the original procedure described by Wu *et al*. (2012). The *E. coli* codon-optimized gene for Arabidopsis UVR8 was introduced into the pET11a expression vector generating a construct carrying an N-terminal 6×His-tag (Genscript). The construct was verified by DNA-sequencing and transformed into the *E. coli* expression host strain BL21. For the recombinant production of functional UVR8, the N-terminal 6×His-tagged UVR8 was overexpressed overnight at 18 °C using 0.2 mM β-d-thiogalactopyranoside for induction (Wu *et al*., 2012). Following the overnight induction, cells were harvested by centrifugation and flash-frozen in liquid nitrogen after which cell pellets were stored at −80 °C. Upon purification of recombinant UVR8, cells were lysed by sonication and soluble protein was separated from insoluble fractions by centrifugation. Further isolation of His-tagged UVR8 was accomplished by immobilized metal-affinity chromatography using a two segmented linear gradient of imidazole. Eluted fractions containing UVR8 was further purified to homogeneity using size-exclusion chromatography as verified by SDS-PAGE. Functionality was verified by electrophoresis assay and fluorescence quenching as described by Wu *et al*. (2012). The UV/Vis absorbance spectrum of UVR8 protein was measured by a standard protocol using a spectrophotometer (Shimadzu UV-1800). To improve the signal-to-noise ratio for spectral regions covering both long and short wavelengths, two separate data sets were recorded for two concentrations of UVR8 protein, 57.0 µM and 4.5 µM, respectively. A 25 mM Tris (pH8), 150 mM NaCl buffer solution supplemented with 1 mM Beta-mercaptoethanol was used. Protein concentrations were determined using a theoretical absorption coefficient of 91900 M^−1^cm^−1^ at 280 nm as determined by the ProtParam data server available through the SIB Swiss Institute of Bioinformatics (Gasteiger *et al*., 2005).

### 2.7 In vitro monomerization of purified UVR8 protein

His-tagged *Arabidopsis thaliana* full-length UVR8 was produced in *Nicotiana benthamiana* after *Agrobacterium tumefaciens* transfection using the pEAQ-HT plasmid (Sainsbury *et al*., 2009) as expression vector. Purification of UVR8 was accomplished using Ni-NTA immobilized metal-affinity and size-exclusion chromatography. Purified UVR8 protein was exposed to UV wavelengths in a 50 µl cuvette using a pulsed Opolette 355+UV tunable laser (Opotek Inc. USA) with a thermostatic cuvette holder at 4 °C, as described in Díaz-Ramos *et al*. (2018), using doses between 1.9 and 6.2 µmol. After exposure, samples were added to 4×SDS sample buffer (250 mM Tris-HCl pH 6.8, 2% (w/v) SDS, 20% (v/v) β-mercaptoethanol, 40% (v/v) glycerol, 0.5% (w/v) bromophenol blue) (O’Hara & Jenkins, 2012) and were subsequently analyzed by SDS-PAGE without boiling (Rizzini *et al.*, 2011). Gels were stained with Coomassie blue to visualize the dimer and monomer bands.

## 3 RESULTS

### 3.1 Gene expression mediated by UVR8 and CRYs after 6 h of sunlight exposure

To get full assessment of changes in transcript abundances induced by sunlight, we performed RNA-seq from samples collected after 6 h of exposure of plants to filtered sunlight. RNA-seq libraries from the same genotype and treatment clustered together showing consistency among biological replicates (Figure S3). Furthermore, we validated the expression profiles obtained by RNA-seq with qRT-PCR using 11 genes responsive to UV radiation, and/or blue light, or involved in hormone responses (Table S1). The high positive correlation, *R*^2^ = 0.85, between the two methods validates the RNA-seq data (Figure S4).

As shown in the Venn diagrams (Figure 2a), in wild type L*er*, out of 3741 DEGs, 2786 responded to blue light (868 UP, 1918 DOWN), 960 to UV-A (406 UP, 554 DOWN) and 653 to UV-B (343 UP, 310 DOWN). Only 101 DEGs responded to all three solar wavebands UV-B, UV-A and blue. Moreover, less than half of the DEGs responding to UV-B were specific, as the remaining ones responded also to UV-A, blue or both. In contrast, of the DEGs responding to UV-A or blue, more than half were specific (Figure 2a).

**Figure 2.**
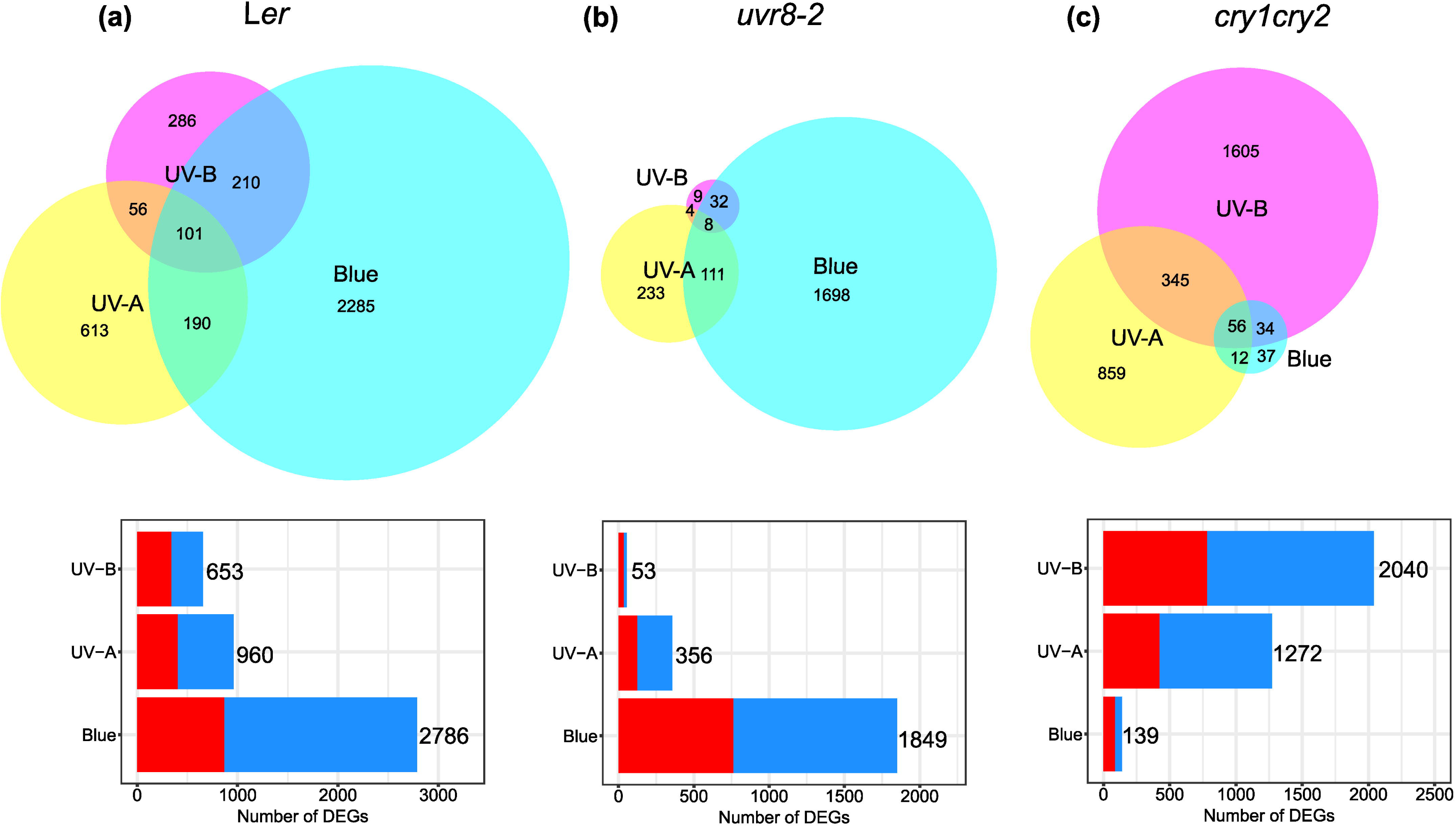
Number of genes differentially expressed in response to 6 h of solar UV-B, UV-A and blue radiation in (a) L*er*, (b) *uvr8-2*, (c) *cry1cry2*. The Venn diagrams show the unique genes for each waveband contrast and the overlap of genes between the waveband contrasts in each genotype. The stacked bar plots show total number of genes responding to the waveband contrasts in each genotype. The red bar refers to genes with increased expression and the blue bar refers to genes with decreased expression. See Figure 1 for the contrasts used to assess the effects of UV-B, UV-A and blue radiation. FC > 1.5 and *P*_adjust_ < 0.05.

In *uvr8-2*, out of 2095 DEGs only 53 (35 UP, 18 DOWN) responded to UV-B (Figure 2b). The number of DEGs responding to UV-A was only 1/3 of that in L*er* (356 differentially expressed, DE; 125 UP, 231 DOWN) while also the number of DEGs responding to blue light was only 2/3 of that in L*er* (1849 DE; 763 UP, 1086 DOWN) (Figure 2a,b). Furthermore, 1/3 of the DEGs responding to blue in *uvr8-2* were unique and not shared by L*er* and *cry1cry2* (Figure S5). Thus, the results confirm our hypothesis that UVR8 plays a role in UV-A perception and also indicate that functional UVR8 modulates gene expression responses to solar blue wavelengths.

In *cry1cry2*, out of 2948 DEGs only 139 (87 UP, 52 DOWN) responded to solar blue light (Figure 2c). Surprisingly for this mutant, out of the 2948 DEGs, 1272 (425 UP, 847 DOWN) still responded to UV-A (Figure 2c). Also, the number of DEGs responding to UV-B increased from 653 in L*er* to 2040 (784 UP, 1256 DOWN) in *cry1cry2* (Figure 2a,c).

To explore the UV-A signaling roles of UVR8 and CRYs in more detail, we next assessed the effects of longer and shorter wavelength regions within UV-A by comparing responses between pairs of filters: UV-A_sw_ (>315 nm vs >350 nm) and UV-A_lw_ (>350 nm vs >400 nm) (Figure 1). In L*er*, out of 190 DEGs, 166 responded to UV-A_sw_ (113 UP, 53 DOWN) and only 26 to UV-A_lw_ (16 UP, 10 DOWN), with only 2 DEGs shared between the two treatments (Figure 3a). In *uvr8-2*, out of 77 DEGs, 7 (3 UP, 4 DOWN) responded to UV-A_sw_, while 72 (17 UP, 55 DOWN) to UV-A_lw_ (Figure 3b). In *cry1cry2*, out of 1057 DEGs, 1050 (340 UP, 710 DOWN) responded to UV-A_sw_ and only 16 (8 UP, 8 DOWN) to UV-A_lw_ (Figure 3c). The number of DEGs responding to UV-A_sw_ in *cry1cry2* was more than six times those responding to UV-A_sw_ in L*er*, a similar but stronger effect to that of UV-B (Figure 3a,c). Furthermore, 3/4 and 9/10 of the DEGs responding to UV-B and UV-A_sw_, respectively, in *cry1cry2* were unique and not shared by L*er* or *uvr8-2* (Figure S5). Here, it should be noted that the individual numbers of DEGs under UV-A_sw_ and UV-A_lw_ add up to a smaller number than the number of DEGs for whole UV-A region (315–400 nm) (cf. Figures 3 and 2). This difference mainly arises from statistically comparing pairs of treatments that correspond to smaller (UV-A_sw_ and UV-A_lw_) or larger (whole UV-A) amounts of sunlight attenuation.

**Figure 3.**
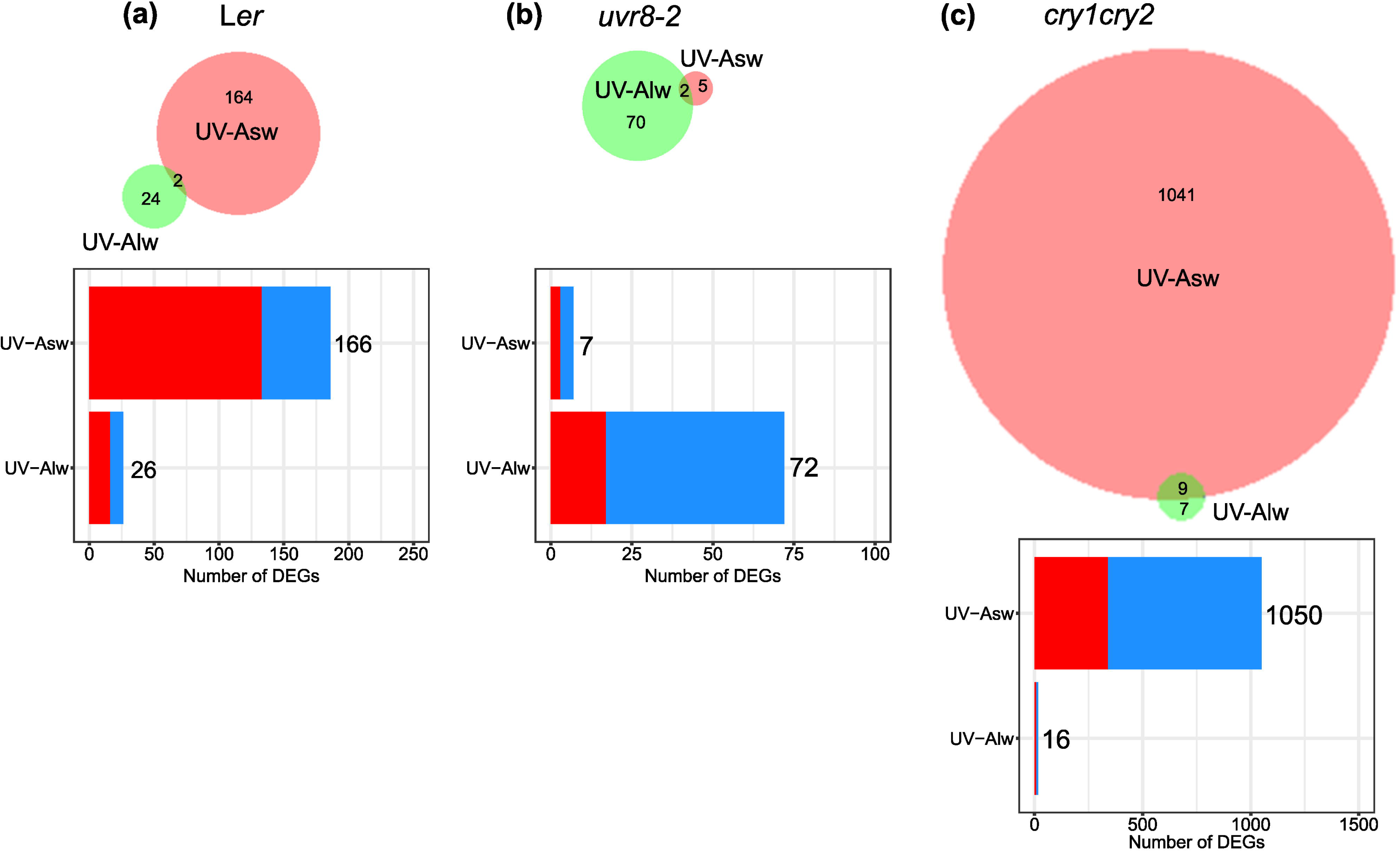
Number of genes differentially expressed in response to 6 h of solar UV-A_sw_ and UV-A_lw_ radiation in (a) L*er*, (b) *uvr8-2*, (c) *cry1cry2*. The Venn diagrams show the unique genes for each waveband contrast and the overlap of genes between the waveband contrasts in each genotype. The stacked bar plots show total number of genes responding to the waveband contrasts in each genotype. The red bar refers to genes with increased expression and the blue bar refers to genes with decreased expression. See Figure 1 for the contrasts used to assess the effects of UV-A_sw_ and UV-A_lw_ radiation. FC > 1.5 and *P*_adjust_ < 0.05.

We next tested whether the requirement of UVR8 vs CRYs observed within the UV-A remained valid when including wavelengths in the UV-B and blue bands in the analysis. For this test, DEGs responding to λ < 350 nm (contrast between >290 nm vs >350 nm) and λ > 350 nm (contrast between >350 nm vs >500 nm) were quantified (Figure 4). The number of DEGs responding to λ < 350 nm in *uvr8-2* drastically decreased to 1/37 of those in L*er* while the number of those responding in *cry1cry2* increased to 2.7 times of those in L*er*. The number of DEGs responding to λ > 350 nm in *cry1cry2* decreased to 1/35 of those in L*er*, while the number of those responding in *uvr8-2* also decreased but only to 4/5 of those in L*er* (Figure 4). This test indicates that in sunlight functional UVR8 is required for transcriptome-wide response to λ < 350 nm and CRYs are required for those to λ > 350 nm and that functional CRYs antagonize this transcriptome-wide response to λ < 350 nm.

**Figure 4.**
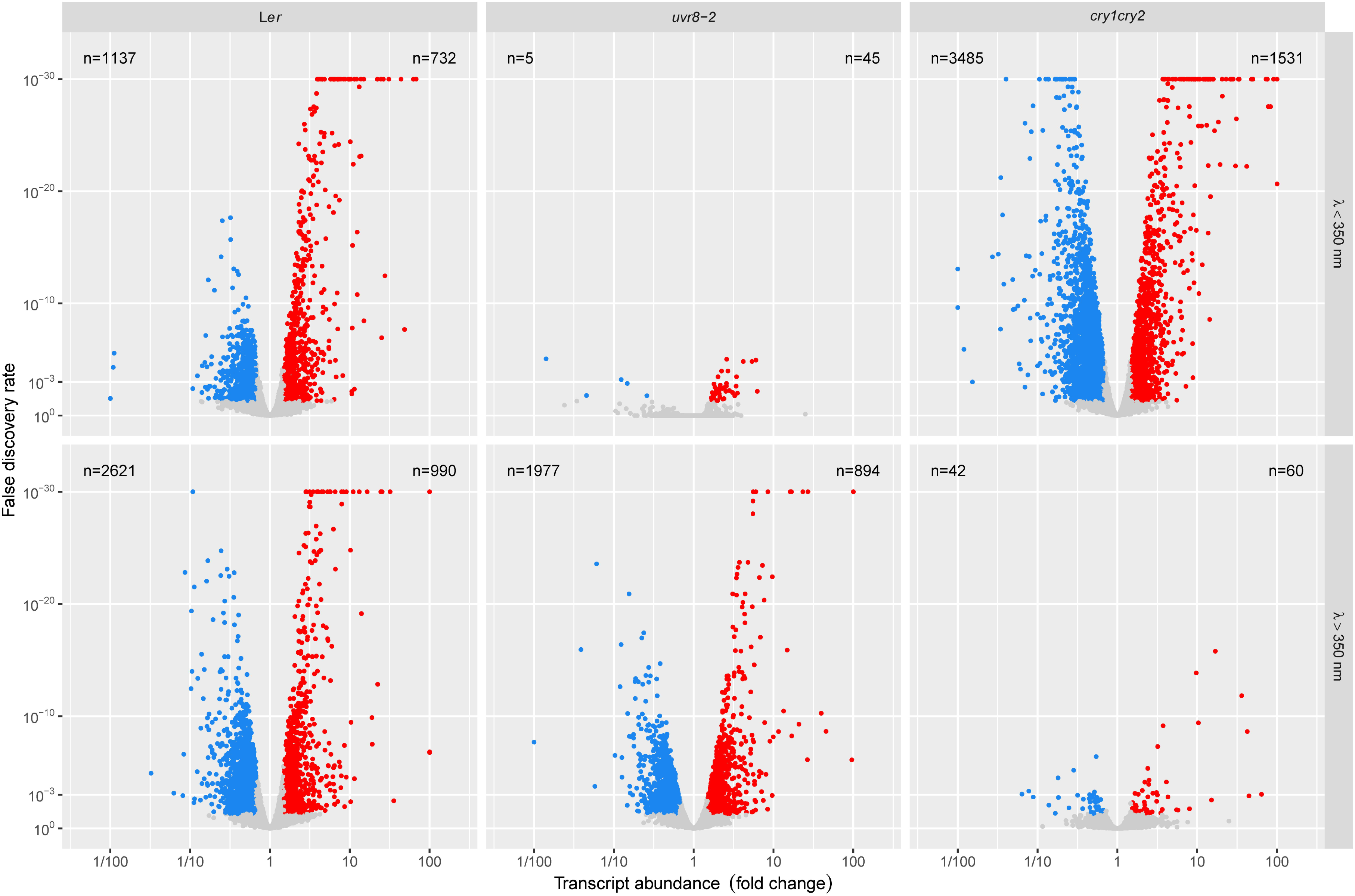
Volcano plots showing DEGs with significantly increased expression (in red), DEGs with significantly decreased expression (in blue) and not significant (in grey) in response to 6 h of solar radiation of λ < 350 nm (290–350 nm) and λ > 350 nm (350–500 nm), λ refers to wavelength, n refers to number of differentially expressed genes. See Figure 1 for the contrasts used to assess the effects of λ < 350 nm and λ > 350 nm. FC > 1.5 and *P*_adjust_ < 0.05.

Our lists of DEGs for the12 waveband-contrast × genotype combinations were enriched for 45 KEGG metabolic pathways in total (Figure S6a-c). The analysis showed that UVR8 mediated the expression of genes involved in all enriched metabolic processes induced by UV-B and UV-A_sw_ in L*er* and *cry1cry2*. Since the flavonoid biosynthesis pathway was still enriched in *uvr8-2* by UV-B, dependence on UVR8 was partial (Figure S6a,b). Expression of genes involved in ribosome biogenesis, protein processing and endocytosis under solar blue required functional UVR8 (Figure S6a,b). CRYs mediated the expression of genes involved in most metabolic pathways regulated by blue light in L*er* and *uvr8-2*, as these responses were missing in *cry1cry2*. The cases where the response to blue light was not fully dependent on CRYs were flavonoid biosynthesis, diterpenoid biosynthesis, phenylalanine metabolism, circadian rhythm, vitamin B6 metabolism, phenylpropanoid biosynthesis. Strikingly, many pathways including photosynthesis, glucosinolates biosynthesis, and plant hormone signal transduction were over-represented in *cry1cry2* compared to L*er* in response to UV-B and UV-A_sw_ (Figure S6a,c).

### 3.2 Photon absorption and monomerization of UVR8

To evaluate whether the actions of UVR8 and CRYs at λ < 350 nm and λ > 350 nm are consistent with their absorption spectra, we estimated the relative number of photons absorbed by both photoreceptors given the solar spectral irradiance on the sampling day under each filter (Figure 5). For UVR8 we used a newly measured absorption spectrum extending into the visible region (Figure S7), while for CRYs we used a published absorption spectrum for CRY2 (Banerjee *et al*., 2007). As a result of the shape of the solar spectrum, the estimates showed that UV-A_sw_ was the band where UVR8 was predicted to absorb most photons. Relative to these maxima, UVR8 was predicted to be the main UV-A_sw_ photoreceptor. In the UV-A_lw_ our estimates showed that both CRYs and UVR8 absorbed a large number of photons (Figure 5). While UVR8 absorbed a considerable number of blue photons, CRY2 absorbed very few UV-B photons.

**Figure 5.**
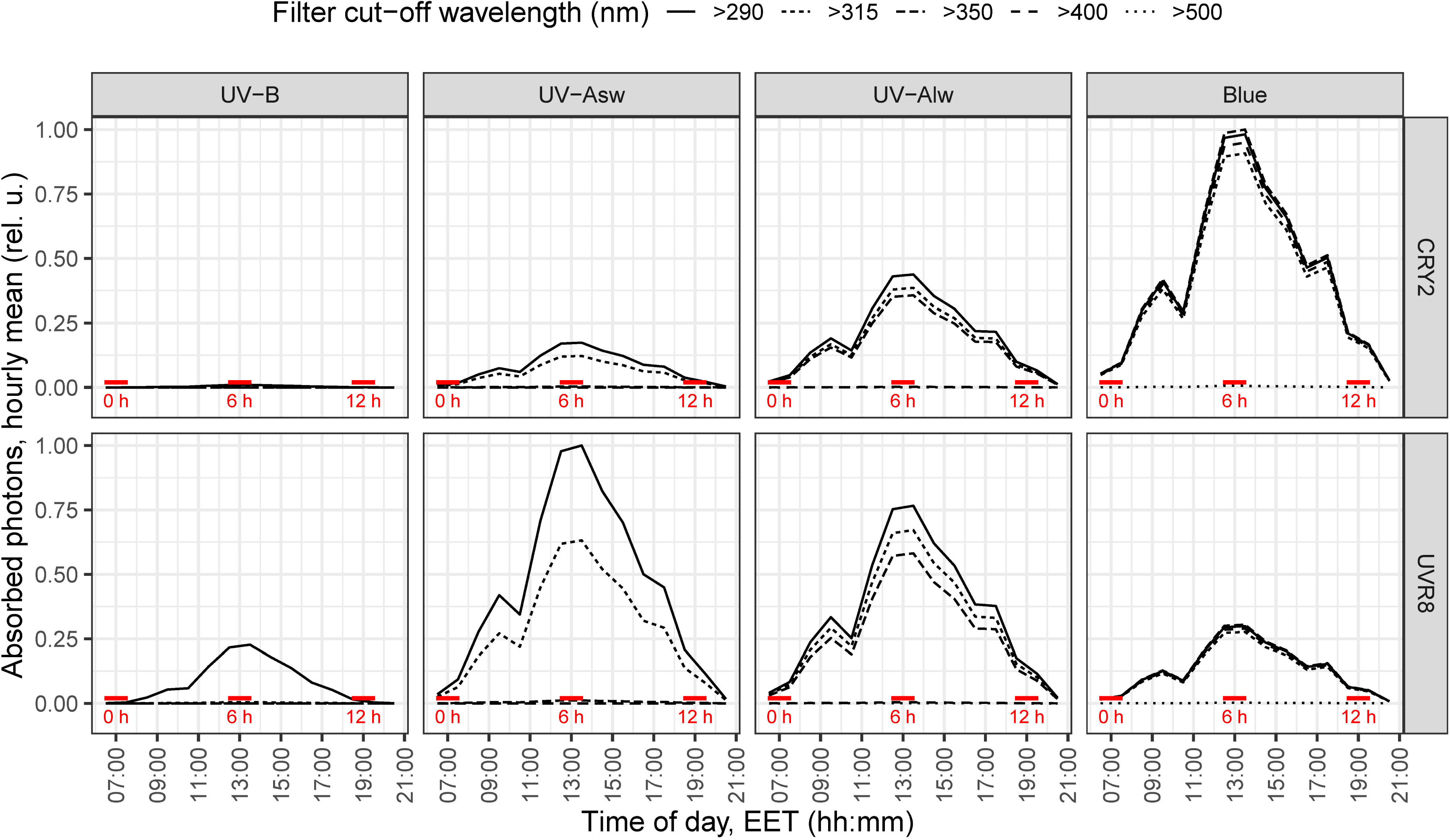
Estimates of solar UV-B, UV-A_sw_, UV-A_lw_ and blue photons absorbed by UVR8 and CRYs proteins throughout the day under the different filters used, expressed relative to their respective daily maximum. The red horizontal lines show when the plants were moved outdoors (0 h) or sampled (6 h and 12 h).

As monomerization of UVR8 is required for UV-B signaling (Rizzini *et al*., 2011), we assessed *in vitro* if laser radiation of different wavelengths within the UV-B and UV-A bands can monomerize purified UVR8 protein. We found that UVR8 monomerized in response to UV wavelengths in the range 300–335 nm but not in 340–350 nm (Figure 6).

**Figure 6.**
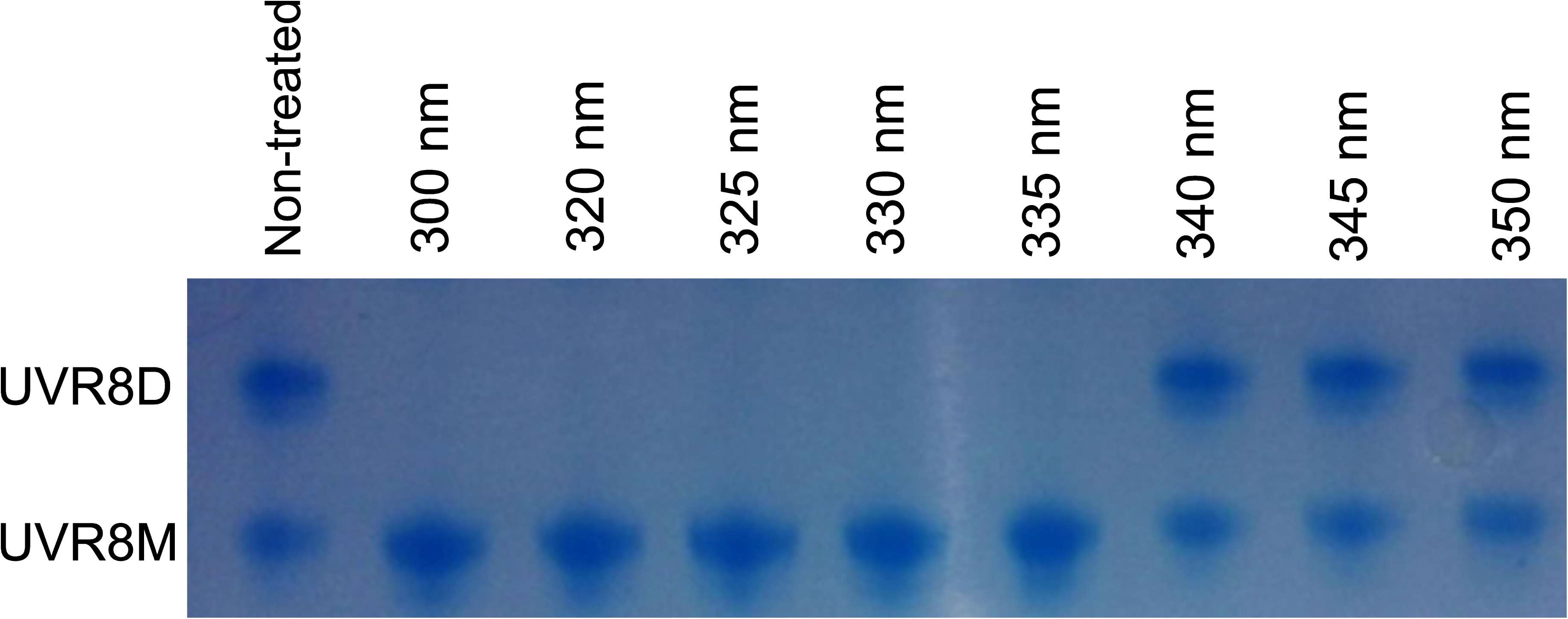
*In vitro* monomerization of purified UVR8 protein from *Nicotiana benthamiana* exposed to UV radiation from a tunable laser. The time of exposure and photon doses at different wavelengths were: 10 min and 1.9 µmol at 300 nm, 30 min and 5.1 µmol at 320 nm, 30 min and 4.5 µmol at 325 nm, 30 min and 3.8 µmol at 330 nm, 30 min and 3.4 µmol at 335 nm, 60 min and 6.2 µmol at 340 nm, 60 min and 5.8 µmol at 345 nm, 60 min and 5.3 µmol at 350 nm. The picture shows the Coomassie stained gel. UVR8D refers to the dimer and UVR8M the monomer. The figure is representative of three repeats.

### 3.3 Cis-motif enrichment

As enrichment of DNA binding motifs can inform about the putative involvement of TFs in the observed gene-expression responses (McLeay & Bailey, 2010), we assessed *in silico* the enrichment of known *cis-*regulatory elements in the promoter regions of the DEGs from each waveband-genotype combination (Figures 7, S8). The analysis identified 187 putative regulatory TFs whose binding motifs were significantly enriched among the DEGs for at least one waveband-genotype combination (Figures 7, S8). The TFs grouped into 12 distinct clusters based on the similarity of their enrichment patterns across four waveband contrasts and three genotypes (Figure 7). Out of these TFs, 53 were themselves differentially expressed. Clusters A, C, D, F, H and I were homogeneous with respect to TF family, whereas the rest were heterogeneous. Within homogeneous clusters several promoter motifs were enriched (Figures 7, S8).

**Figure 7.**
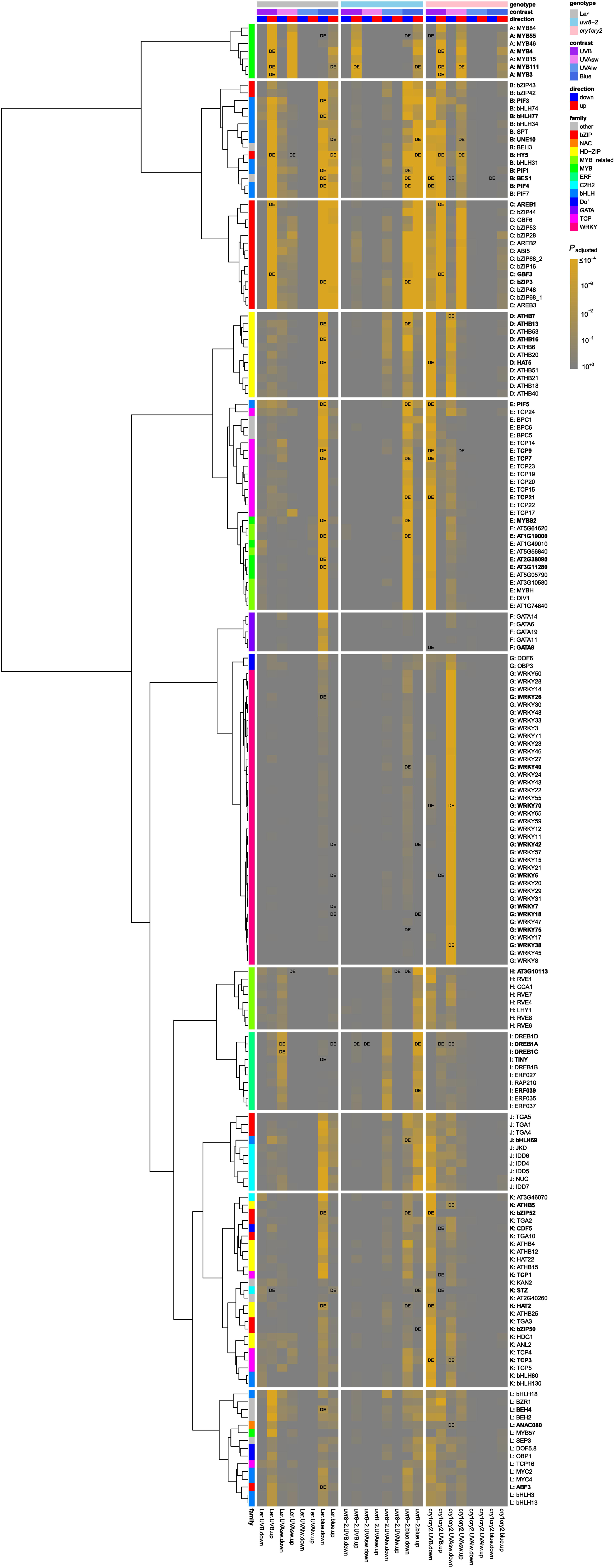
The *in silico* enrichment of DNA-binding motifs in our RNA-seq data showing 187 putative regulatory transcription factors (TFs). See Figure S8 for the position weight matrix of the enriched motifs. The enrichments were done in 1000 base pairs upstream of the coding regions of the DEGs from each waveband contrast and genotype combination. TFs with *P*_adjust_ < 0.01 in at least one contrast and genotype combination were included in the figure. Of the 187 enriched TFs, those which were also differentially expressed in our experiment are presented in bold letters and shown as “DE” in the figure.

In cluster A, several MYB TFs including MYB111 were predicted to regulate the expression of genes with increased transcript abundance in response to UV-B and blue in all genotypes, while in response to UV-A_sw_ significant enrichment was found only in L*er* and *cry1cry2* (cluster A, Figure 7). TFs in this group were the only ones enriched for genes responding to UV-B in *uvr8-2*. Out of these seven TFs, four were themselves differentially expressed in response to the treatments.

HY5, PIF1, PIF3, PIF4, PIF7, BES1 were grouped in cluster B, and were predicted to regulate the expression of genes with increased transcript abundance in response to solar UV-B in L*er* and *cry1cry2* (Figure 7) and of genes with either increased or decreased transcript abundance in response to solar blue in L*er* and *uvr8-2*. Most of these same TFs were enriched for two additional responses only in *cry1cry2*: decreased transcript abundance by UV-B and increased transcript abundance by UV-A_sw_. Out of these 15 TFs, seven, including HY5, PIF1, PIF3, PIF4 and BES1, were themselves differentially expressed. Cluster C which follows a response pattern similar to that of cluster B groups 14 bZIP TFs of which three were differentially expressed.

Eleven members of HD-ZIP TFs including HAT5 grouped in cluster D and were enriched for decreased transcript abundance only in response to blue light in L*er* and *uvr8-2* but in response to UV-B and UV-A_sw_ only in *cry1cry2* (Figure 7). Out of these 11 TFs, four including HAT5 were differentially expressed. Cluster E which follows a response pattern similar to that of cluster D contains TFs from a mix of families of which eight were differentially expressed.

Five GATA TFs were enriched for DEGs repressed by blue only in L*er* whereas in the other two mutants these TFs were not enriched for any response (cluster F, Figure 7). WRKY TFs were enriched only for DEGs with decreased expression in response to UV-A_sw_ in *cry1cry2* (cluster G). The number of differentially expressed TFs were 1 and 9 in clusters F and G, respectively. The remaining clusters (H-L) showed multiple but poorly defined patterns of enrichment.

### 3.4 Transcript abundance after 6 h and 12 h

We used qRT-PCR to determine changes in transcript abundance after 6 h (mid-day) and after 12 h (before sunset) of exposure to filtered solar radiation, allowing the assessment of transcript abundance at two times of the day when solar UV-B:UV-A photon ratio was very different. We tested 11 genes (Figures 8, S9). The three-way interaction, treatment × genotype × exposure time, was significant for four genes: *RUP2* (involved in UVR8 signaling), *CHALCONE SYNTHASE* (*CHS*) and *CHALCONE ISOMERASE* (*CHI*) (flavonoid biosynthesis); and *SOLANESYL DIPHOSPHATE SYNTHASE 1* (*SPS1*) (ubiquinone biosynthesis) (Figure 8, Table S2). This indicates that in sunlight the role of UVR8 and CRYs in the regulation of transcript abundance of these genes changed in time. For these four genes across all genotypes, the response to filter treatments at 6 h was stronger than at 12 h.

**Figure 8.**
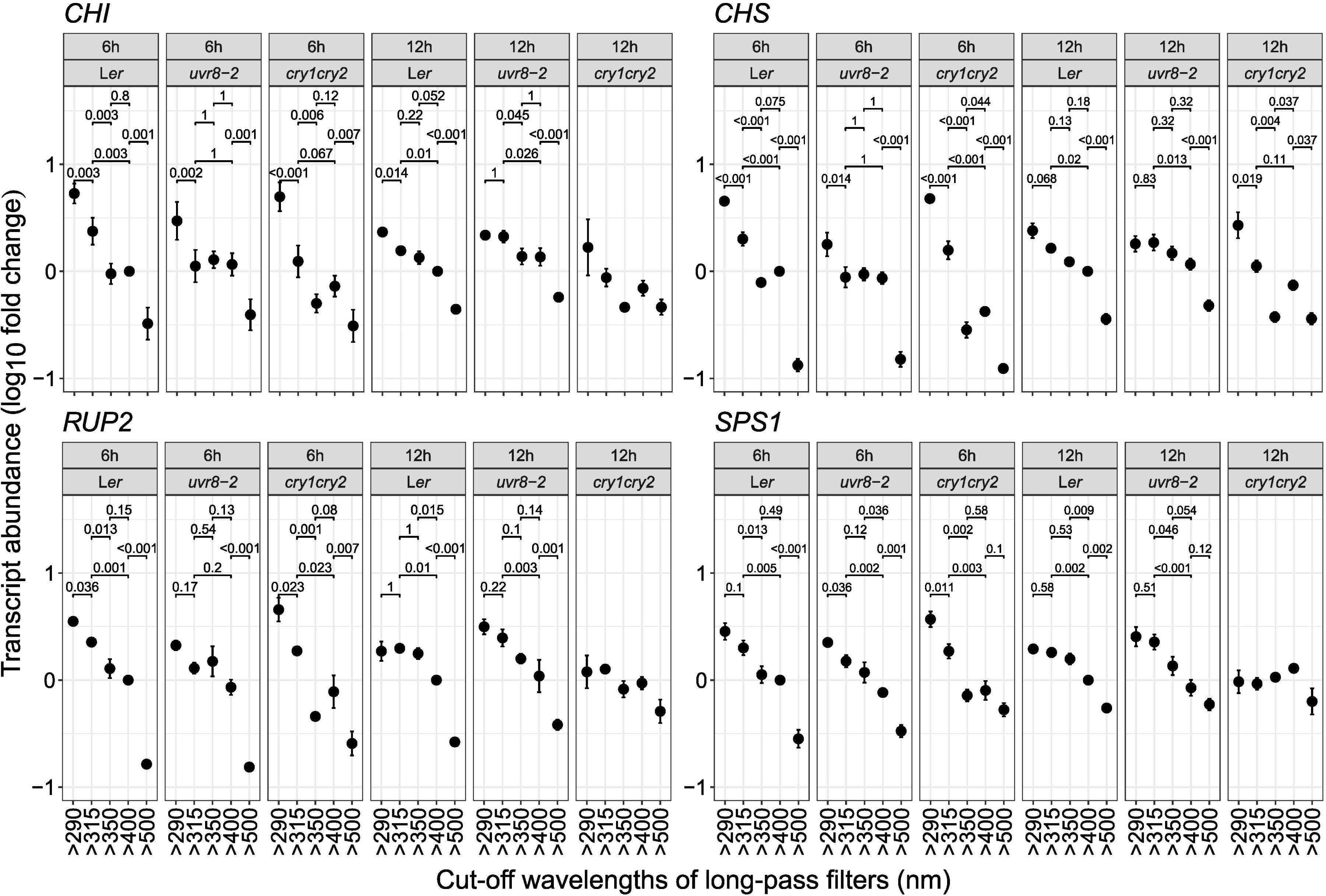
Transcript abundance (log_10_FC) of four genes *CHI*, *CHS*, *RUP2* and *SPS1* after 6 h and 12 h of treatment outdoors. These genes showed a significant triple interaction (genotype × radiation treatment × exposure time). Mean ±1SE from four biological replicates. The horizontal bars show *P*_adjust_ values for pair-wise comparisons between treatments within each genotype. *P*_adjust_ values for pair-wise contrasts are shown only in those panels where the overall effect of filter treatment within a genotype and time point was significant (see Table S2).

Solar UV-B at 6 h increased transcript abundance of *CHI* and *CHS* in all genotypes and of *RUP2* in L*er* and *cry1cry2* but not in *uvr8-2* (Figure 8). This response to UV-B was stronger in *cry1cry2* than in L*er* for all these genes. Solar UV-A_sw_ at 6 h increased transcript abundance of *CHI*, *CHS*, *RUP2* and *SPS1* in L*er* and *cry1cry2* but not in *uvr8-2*. As for UV-B, the transcript abundance response to 6 h of UV-A_sw_ was stronger in *cry1cry2* than in L*er* for these four genes. Both solar UV-B and UV-A_sw_ at 12 h increased the abundance of *CHS*, but only in *cry1cry2*. These results support regulation of transcript abundance by UVR8 in both UV-B and UV-A_sw_, antagonized by CRYs.

Solar UV-A_lw_ at both 6 h and 12 h decreased the transcript abundance of *CHS* in *cry1cry2* (Figure 8). UV-A_lw_ at 12 h increased the transcript abundance of *RUP2* in L*er* and of *SPS1* in both L*er* and *uvr8-2* but not in *cry1cry2*. Overall, transcript abundance was less responsive to UV-A_lw_ than to UV-A_sw_. Solar blue light at 6 h increased transcript abundance of *CHI*, *CHS* and *RUP2* in all genotypes whereas of *SPS1* in L*er* and *uvr8-2* but not in *cry1cry2*. Blue light at 12 h increased the abundance of *CHI* and *RUP2* only in L*er* and *uvr8-2*, and of *CHS* in all three genotypes, but less in *cry1cry2*. These results support regulation of transcript abundance by CRYs in UV-A_lw_ and blue light.

## 4 DISCUSSION

### 4.1 Effective range of wavelengths for action of UVR8 and CRYs in sunlight

In plant photobiology the consensus has been that UVR8 and CRYs function as UV-B and blue/UV-A photoreceptors, respectively (Ahmad & Cashmore, 1993; Rizzini *et al*., 2011). However, here we show that UVR8 mediates transcriptome-wide changes in response to both solar UV-A_sw_ and UV-B (Figures 2–4). Assuming that UVR8 monomers are required for signaling and response (Rizzini *et al*., 2011), for UVR8 to mediate responses to UV-A_sw_ it must absorb enough photons at these longer wavelengths and monomerize. Our *in silico* estimates based on spectral absorbance predict that UVR8 absorbs more UV-A_sw_ photons than UV-B photons in sunlight (Figure 5), because sunlight contains at least 30 times more UV-A_sw_ photons than UV-B photons (Aphalo, 2018). This explains why the role of UVR8 in UV-A_sw_ perception has not been observed in earlier studies using artificial light with unrealistically high UV-B:PAR and low UV-A:PAR ratios. We also show that *in vitro* UVR8 dimers convert to monomers when exposed to radiation of wavelengths between 300 nm and 335 nm but not in response to longer wavelengths (Figure 6). As the UVR8 protein does absorb photons at wavelengths longer than 335 nm, a possible explanation for this transition between 335 nm and 340 nm is a threshold in the energy per photon required for monomerization. The dose we used at 335 nm was more than 3000 times the maximum used by Díaz-Ramos *et al*. (2018), who observed *in vitro* almost complete monomerization at 310 nm, the longest wavelength they investigated.

We also found that UVR8 affected blue light-induced gene expression, as fewer and in part different genes responded to blue light in *uvr8-2* when compared to L*er* (Figures 2a,b, S5). This effect was unexpected, as the effect of blue light was assessed in a background of strongly attenuated UV-B and UV-A. Furthermore, KEGG pathway analysis indicates that functional UVR8 might be required for blue-light-dependent expression of genes involved in ribosome biogenesis, protein processing and endocytosis (Figures S6a-c). Our estimates also predict that photon absorption by UVR8 in sunlight extends as far as the blue region (Figure S7), where we observed UVR8-dependent modulation of these specific responses to solar blue instead of a clear-cut requirement as at shorter wavelengths. Although evidence for perception of blue light by UVR8 is weak, the lack of monomerization in response to wavelengths longer than 335 nm suggests that photoreception by UVR8 at longer wavelengths would have to depend on a different mechanism.

Our transcriptomic data at 6 h (solar noon) indicate that CRYs are the main photoreceptors mediating gene expression responses to solar blue and UV-A_lw_, but not to UV-A_sw_. This contrasts with the currently accepted role of CRYs in perception of the whole UV-A waveband (Yu *et al*., 2010). Yet, the absence of CRYs increased the number of DEGs up to three times in response to UV-B and up to six times in response to UV-A_sw_ (Figures 2a,c and 3a,c), an unexpectedly large effect affecting many metabolic processes (Figure S6). However, while CRYs absorb comparatively fewer photons at these shorter wavelengths than in the blue, the effect of UV-B and UV-A_sw_ exposure was assessed in a background of UV-A_lw_ and blue radiation, a condition under which CRY signaling was activated in the WT but not in *cry1cry2*. Thus, the previously described negative regulation of four UVR8-mediated genes by CRYs in UV-B and UV-A_sw_ (Rai *et al*., 2019) was now expanded to the whole transcriptome indicating an interaction between UVR8 and CRYs leading to wide-ranging regulation of primary and secondary metabolism (Figures 2–4, S6). Furthermore, this negative regulation was observed using qRT-PCR at both 6 h and 12 h for *CHS* (Figure 8) indicating that this effect can persist until the end of the photoperiod even though UV-B irradiance was very low at this time, suggesting a carry-over effect. The transcript abundance of *RUP1* and *RUP2* was increased in response to UV-A_sw_ in *cry1cry2* compared to L*er* (Figure 8), indicating that the crosstalk between UV-B/UV-A_sw_ and UV-A_lw_/blue signaling pathways may involve RUPs. In addition, a recent study demonstrated that both UVR8 and CRYs use VP motifs to compete for binding the WD40 domain of COP1 (Lau *et al*., 2019). Therefore, crosstalk between the two signaling pathways could involve COP1. These earlier results together with our new observations provide a good starting point for future studies on the molecular mechanism of interaction between UVR8 and CRYs in sunlight and its relevance to plant adaptation and acclimation to diurnal and seasonal variation in the solar spectrum.

### 4.2 Putative TFs behind different patterns of gene expression response

Our promoter enrichment analysis highlights the possible roles of several TFs in controlling the observed gene expression responses downstream of UVR8 and CRYs. The analysis predicted that MYB TFs, known regulators of flavonol accumulation (Stracke *et al*., 2007), regulate the expression of genes responding to UV-B partially independent of UVR8, and of those responding to UV-A_sw_ through UVR8 (Figure 7, cluster A). The data also show that HAT5 and PIF5 (clusters D, E) are predicted to specifically regulate gene expression in response to solar blue through CRYs. These results in sunlight agree with a previous report where PIF5 is shown to function downstream of CRYs to mediate hypocotyl elongation in response to artificial blue light (Pedmale *et al*., 2016).

Although earlier work has emphasized the role of HY5 as a master TF central to responses to UV radiation and blue light (Brown & Jenkins, 2008; Favory *et al*., 2009; Gangappa & Botto, 2016), the array of response patterns of transcript abundance and motif enrichment observed here indicate that several TFs play key roles in downstream signaling leading to gene expression. In addition to HY5, the known regulators of UV and blue light signaling and photomorphogenesis including PIF1, PIF3, PIF4, PIF7 and BES1 (Hayes *et al*., 2014; Gangappa & Botto, 2016; Pedmale *et al*., 2016; Liang *et al*., 2018; Wang *et al*., 2018) were predicted to regulate gene expression in response to solar UV-B through UVR8, and in response to solar blue through CRYs (Figure 7, cluster B). Our data also indicate that HY5, PIFs, BES1, HAT5, WRKYs and many other TFs could regulate the expression of genes responsive to UV-B or UV-A_sw_ and require both UVR8 and CRYs (Figure 7, clusters B, C, D and G). This shows that both UVR8 and CRY signaling employ some of the same TFs for gene expression. However, the multiple points of interaction for crosstalk between UV-B, UV-A_sw,_ UV-A_lw_ and blue light signaling pathways downstream of UVR8, CRYs and other photoreceptors remain to be explored.

### 4.3 Implications and conclusions

With few exceptions, gene expression in response to solar UV-B and UV-A_sw_ depended on UVR8, while that in response to UV-A_lw_ and blue light depended on CRYs. Why the “UV-B photoreceptor” UVR8 played a role in the perception of solar UV-A_sw_ can be explained by the numbers of solar UV-B and UV-A_sw_ photons predicted to be absorbed by UVR8, a physico-chemical mechanism. Our prediction of photons absorbed by UVR8 was made possible by the new spectral absorbance data we report, demonstrating the usefulness of extending such measurements far along the tails of absorption spectra. We also observed *in vitro* monomerization of UVR8 dimers exposed to wavelengths between 300 and 335 nm but not when exposed to longer ones, extending previous knowledge into longer wavelengths. This lack of monomerization may explain why UVR8 does not play an important role in the perception of UV-A_lw_. Thus, we describe the mechanism by which the steep slope of the solar spectrum in the UV region shifts perception of solar radiation by UVR8 towards longer wavelengths than frequently assumed.

When considering both UVR8 and CRYs, we observed that the transcriptome-wide response triggered by UV-B and UV-A_sw_ exposure was very strongly and negatively regulated by CRYs. The reverse effect, modulation by UVR8 of gene expression in response to blue light was also observed although it was much weaker. These results demonstrate for the first time the extent of the effect of interactions downstream of UVR8 and CRYs on the transcriptome. These data also allowed us to putatively identify several metabolic pathways affected by the interaction.

Specific groups of TFs were predicted *in silico* to control cascades of gene expression corresponding to different patterns of transcriptome response to wavelengths across genotypes, patterns which can only arise as the result of a complex signaling network including multiple points of interaction downstream of UVR8 and CRYs. This prediction highlights that current models of signaling downstream of UVR8 and CRYs, rather unsurprisingly, describe only the top portion of a much deeper and ramified signaling network. As our study demonstrates, experiments combining the use of multiple light treatments and multiple mutants in a factorial design allow teasing out some of the signaling complexity that is missing from current models. The *in-silico* predictions we report can guide the development of hypotheses about the mechanisms and players involved in signaling, hypotheses that will need to be tested in future experiments.

As the wavelength boundaries for effective sensitivity of UVR8- and CRY-mediated perception of sunlight do not coincide with the definitions of UV-B and UV-A radiation in common use (Björn, 2015), we consider that quantification of solar radiation based on these definitions is only marginally useful when studying sunlight perception by plants, i.e., photomorphogenesis rather than stress damage. Even more important, is that in both irradiation- and waveband-attenuation experiments different regions within UV-A will trigger responses through different photoreceptors, possibly resulting in contradictory or confusing results. In the present study, splitting the UV-A waveband at 350 nm into UV-A_sw_ and UV-A_lw_ was the key to revealing the effective roles of UVR8 and CRYs in the perception of UV radiation in sunlight. Thus, as we routinely do for red and far-red light in the visible, it is also very profitable to use plant-photomorphogenesis-specific waveband definitions to characterize radiation in the UV-A region.

## ACKNOWLEDGEMENTS

We acknowledge Petri Auvinen (University of Helsinki) for RNA-seq. Funding by Academy of Finland (252548) to PJA, and (307335) to MB and JS; EDUFI Fellowship, Finnish Cultural Foundation and Doctoral Program in Plant Sciences funding (University of Helsinki) to NR; Knowledge foundation (20130164) and Swedish Research Council Formas (942-2015-516) to ÅS; Strategic Young Researchers Recruitment Programme (Örebro University) to LOM.

## 6 AUTHORSHIP

PJA and LOM planned the research. NR, MB, ÅS, PJA, and LOM designed experiments. NR, AO’H, DF, KR, FW, AVL, and LOM performed experiments. NR, OS, JS, PJA, and LOM analyzed data. NR, PJA and LOM wrote the paper with contributions from MB, JS, ÅS, GIJ, and TL. All authors commented and approved the manuscript. PJA and LOM contributed equally as senior authors.

## References

Ahmad M., Cashmore A.R. (1993). HY4 gene of *A. thaliana* encodes a protein with characteristics of a blue-light photoreceptor. Nature, 366, 162–166.

Ambrosini G., Groux R., Bucher P. (2018). PWMScan: a fast tool for scanning entire genomes with a position-specific weight matrix (J Hancock, Ed.). Bioinformatics, 34, 2483–2484.

Andrews S. (2014). FastQC A quality control tool for high throughput sequence data. Babraham Bioinformatics. URL https://www.bioinformatics.babraham.ac.uk/projects/fastqc/ [accessed 12 June 2019].

Aphalo P.J. (2015). The r4photobiology suite: spectral irradiance. UV4Plants Bulletin, 2015, 21–29.

Aphalo P.J. (2018). Exploring temporal and latitudinal variation in the solar spectrum at ground level with the TUV model. UV4Plants Bulletin, 2018, 45–56.

Banerjee R., Schleicher E., Meier S., Viana R.M., Pokorny R., Ahmad M., Bittl R., Batschauer A. (2007). The signaling state of Arabidopsis cryptochrome 2 contains flavin semiquinone. The Journal of Biological Chemistry, 282, 14916–14922.

Björn L.O. (2015). History Ultraviolet-A, B, and C. UV4Plants Bulletin, 2015, 17–18.

Blomster T., Salojärvi J., Sipari N., Brosché M., Ahlfors R., Keinänen M., Overmyer K., Kangasjärvi J. (2011). Apoplastic reactive oxygen species transiently decrease auxin signaling and cause stress-induced morphogenic response in Arabidopsis. Plant Physiology, 157, 1866–1883.

Bolger A.M., Lohse M., Usadel B. (2014). Trimmomatic: a flexible trimmer for Illumina sequence data. Bioinformatics, 30, 2114–2120.

Bray N.L., Pimentel H., Melsted P., Pachter L. (2016). Near-optimal probabilistic RNA-seq quantification. Nature Biotechnology, 34, 525–527.

Brelsford C.C., Morales L.O., Nezval J., Kotilainen T.K., Hartikainen S.M., Aphalo P.J., Robson T.M. (2018). Do UV-A radiation and blue light during growth prime leaves to cope with acute high light in photoreceptor mutants of *Arabidopsis thaliana*c? Physiologia Plantarum, 165, 537–554.

Brown B.A., Cloix C., Jiang G.H., Kaiserli E., Herzyk P., Kliebenstein D.J., Jenkins G.I. (2005). A UV-B-specific signaling component orchestrates plant UV protection. Proceedings of the National Academy of Sciences of the United States of America, 102, 18225–18230.

Brown B.A., Jenkins G.I. (2008). UV-B signaling pathways with different fluence-rate response profiles are distinguished in mature Arabidopsis leaf tissue by requirement for UVR8, HY5, and HYH. Plant Physiology, 146, 576–588.

Christie J.M., Arvai A.S., Baxter K.J., Heilmann M., Pratt A.J., O’Hara A., Kelly S.M., Hothorn M., Smith B.O., Hitomi K., et al. (2012). Plant UVR8 photoreceptor senses UV-B by tryptophan-mediated disruption of cross-dimer salt bridges. Science, 335, 1492–1496.

Cloix C., Kaiserli E., Heilmann M., Baxter K.J., Brown B.A., O’Hara A., Smith B.O., Christie J.M., Jenkins G.I. (2012). C-terminal region of the UV-B photoreceptor UVR8 initiates signaling through interaction with the COP1 protein. Proceedings of the National Academy of Sciences of the United States of America, 109, 16366–16370.

Díaz-Ramos L.A., O’Hara A., Kanagarajan S., Farkas D., Strid Å., Jenkins G.I. (2018). Difference in the action spectra for UVR8 monomerisation and *HY5* transcript accumulation in Arabidopsis. Photochemical and Photobiological Sciences, 17, 1108–1117.

Emde C., Buras-Schnell R., Kylling A., Mayer B., Gasteiger J., Hamann U., Kylling J., Richter B., Pause C., Dowling T., et al. (2016). The libRadtran software package for radiative transfer calculations (version 2.0.1). Geoscientific Model Development, 9, 1647–1672.

Favory J.-J., Stec A., Gruber H., Rizzini L., Oravecz A., Funk M., Albert A., Cloix C., Jenkins G.I., Oakeley E.J., et al. (2009). Interaction of COP1 and UVR8 regulates UV-B-induced photomorphogenesis and stress acclimation in Arabidopsis. The EMBO Journal, 28, 591–601.

Gangappa S.N., Botto J.F. (2016). The multifaceted roles of HY5 in plant growth and development. Molecular Plant, 9, 1353–1365.

Gasteiger E., Hoogland C., Gattiker A., Duvaud S., Wilkins M.R., Appel R.D., Bairoch A. (2005). Protein identification and analysis tools on the ExPASy server. In The Proteomics Protocols Handbook, pp. 571–607. Humana Press, Totowa, New Jersey.

Gruber H., Heijde M., Heller W., Albert A., Seidlitz H.K., Ulm R. (2010). Negative feedback regulation of UV-B-induced photomorphogenesis and stress acclimation in Arabidopsis. Proceedings of the National Academy of Sciences of the United States of America, 107, 20132–20137.

Hayes S., Velanis C.N., Jenkins G.I., Franklin K.A. (2014). UV-B detected by the UVR8 photoreceptor antagonizes auxin signaling and plant shade avoidance. Proceedings of the National Academy of Sciences of the United States of America, 111, 11894–11899.

Heijde M., Ulm R. (2013). Reversion of the Arabidopsis UV-B photoreceptor UVR8 to the homodimeric ground state. Proceedings of the National Academy of Sciences of the United States of America, 110, 1113–1118.

Holm S. (1979). A simple sequentially rejective multiple test procedure BibSonomy. Scandinavian Journal of Statistics, 6, 65–70.

Huang X., Ouyang X., Yang P., Lau O.S., Chen L., Wei N., Deng X.W. (2013). Conversion from CUL4-based COP1-SPA E3 apparatus to UVR8-COP1-SPA complexes underlies a distinct biochemical function of COP1 under UV-B. Proceedings of the National Academy of Sciences of the United States of America, 110, 16669–16674.

Kaiserli E., Jenkins G.I. (2007). UV-B promotes rapid nuclear translocation of the Arabidopsis UV-B specific signaling component UVR8 and activates its function in the nucleus. The Plant Cell, 19, 2662–2673.

Kallio M.A., Tuimala J.T., Hupponen T., Klemelä P., Gentile M., Scheinin I., Koski M., Käki J., Korpelainen E.I. (2011). Chipster: user-friendly analysis software for microarray and other high-throughput data. BMC Genomics, 12, 507.

Kerr J.B., Fioletov V.E. (2008). Surface ultraviolet radiation. Atmosphere-Ocean, 46, 159–184.

Khan A., Fornes O., Stigliani A., Gheorghe M., Castro-Mondragon J.A., van der Lee R., Bessy A., Chèneby J., Kulkarni S.R., Tan G., et al. (2018). JASPAR 2018: update of the open-access database of transcription factor binding profiles and its web framework. Nucleic Acids Research, 46, D260–D266.

Kleine T., Kindgren P., Benedict C., Hendrickson L., Strand Å. (2007). Genome-wide gene expression analysis reveals a critical role for CRYPTOCHROME1 in the response of Arabidopsis to high irradiance. Plant Physiology, 144, 1391–1406.

Kolde R. (2019). pheatmap: Pretty Heatmaps. R package version 1.0.12. URL https://cran.r-project.org/web/packages/pheatmap/index.html [accessed 12 June 2019].

Lau K., Podolec R., Chappuis R., Ulm R., Hothorn M. (2019). Plant photoreceptors and their signaling components compete for binding to the ubiquitin ligase COP1 using their VP-peptide motifs. The EMBO Journal, 38, e102140.

Legendre P. (2018). lmodel2: Model II Regression. R package version 1.7–3. URL https://CRAN.R-project.org/package=lmodel2 [accessed 12 June 2019].

Lian H.L., He S.B., Zhang Y.C., Zhu D.M., Zhang J.Y., Jia K.P., Sun S.X., Li L., Yang H.Q. (2011). Blue-light-dependent interaction of cryptochrome 1 with SPA1 defines a dynamic signaling mechanism. Genes and Development, 25, 1023–1028.

Liang T., Mei S., Shi C., Yang Y., Peng Y., Ma L., Wang F., Li X., Huang X., Yin Y., et al. (2018). UVR8 interacts with BES1 and BIM1 to regulate transcription and photomorphogenesis in Arabidopsis. Developmental Cell, 44, 512–523.

Lin C. (2000). Plant blue-light receptors. Trends in Plant Science, 5, 337–342.

Lindfors A., Heikkilä A., Kaurola J., Koskela T., Lakkala K. (2009). Reconstruction of solar spectral surface UV irradiances using radiative transfer simulations. Photochemistry and Photobiology, 85, 1233–1239.

Liu H., Liu B., Zhao C., Pepper M., Lin C. (2011a). The action mechanisms of plant cryptochromes. Trends in Plant Science, 16, 684–691.

Liu B., Yang Z., Gomez A., Liu B., Lin C., Oka Y. (2016). Signaling mechanisms of plant cryptochromes in *Arabidopsis thaliana*. Journal of Plant Research, 129, 137–148.

Liu H., Yu X., Li K., Klejnot J., Yang H., Lisiero D., Lin C. (2008). Photoexcited CRY2 interacts with CIB1 to regulate transcription and floral initiation in Arabidopsis. Science, 322, 1535–1539.

Liu B., Zuo Z., Liu H., Liu X., Lin C. (2011b). Arabidopsis cryptochrome 1 interacts with SPA1 to suppress COP1 activity in response to blue light. Genes and Development, 25, 1029–1034.

Mazzella M.A., Cerdán P.D., Staneloni R.J., Casal J.J. (2001). Hierarchical coupling of phytochromes and cryptochromes reconciles stability and light modulation of Arabidopsis development. Development, 128, 2291–2299.

McCarthy D.J., Chen Y., Smyth G.K. (2012). Differential expression analysis of multifactor RNA-Seq experiments with respect to biological variation. Nucleic Acids Research, 40, 4288–4297.

McLeay R.C., Bailey T.L. (2010). Motif Enrichment Analysis: a unified framework and an evaluation on ChIP data. BMC Bioinformatics, 11, 165.

Morales L.O., Brosché M., Vainonen J., Jenkins G.I., Wargent J.J., Sipari N., Strid Å., Lindfors A.V., Tegelberg R., Aphalo P.J. (2013). Multiple roles for UV RESISTANCE LOCUS8 in regulating gene expression and metabolite accumulation in Arabidopsis under solar ultraviolet radiation. Plant Physiology, 161, 744–759.

Neff M.M., Chory J. (1998). Genetic interactions between Phytochrome A, Phytochrome B, and Cryptochrome 1 during Arabidopsis development. Plant Physiology, 118, 27–36.

O’Hara A., Jenkins G.I. (2012). In vivo function of tryptophans in the Arabidopsis UV-B photoreceptor UVR8. The Plant Cell, 24, 3755–3766.

O’Malley R.C., Huang S.-S.C., Song L., Lewsey M.G., Bartlett A., Nery J.R., Galli M., Gallavotti A., Ecker J.R. (2016). Cistrome and epicistrome features shape the regulatory DNA landscape. Cell, 165, 1280–1292.

Ohgishi M., Saji K., Okada K., Sakai T. (2004). Functional analysis of each blue light receptor, cry1, cry2, phot1, and phot2, by using combinatorial multiple mutants in Arabidopsis. Proceedings of the National Academy of Sciences, 101, 2223–2228.

Pedmale U.V., Huang S.C., Zander M., Cole B.J., Hetzel J., Ljung K., Reis P.A.B., Sridevi P., Nito K., Nery J.R., et al. (2016). Cryptochromes interact directly with PIFs to control plant growth in limiting blue light. Cell, 164, 233–245.

Pinheiro J., Bates D., DebRoy S., Sarkar D., R Core Team. (2018). nlme: Linear and Nonlinear Mixed Effects Models. URL https://cran.r-project.org/web/packages/nlme/index.html [accessed 12 June 2019].

Podolec R., Ulm R. (2018). Photoreceptor-mediated regulation of the COP1/SPA E3 ubiquitin ligase. Current Opinion in Plant Biology, 45, 18–25.

R Core Team. (2018). A language and environment for statistical computing. R Foundation for Statistical Computing. URL https://www.r-project.org/ [accessed 12 June 2019].

Rai N., Neugart S., Yan Y., Wang F., Siipola S.M., Lindfors A.V., Winkler J.B., Albert A., Brosché M., Lehto T., et al. 2019. How do cryptochromes and UVR8 interact in natural and simulated sunlight? Journal of Experimental Botany, 70, 4975–4990.

Ritchie M.E., Phipson B., Wu D., Hu Y., Law C.W., Shi W., Smyth G.K. (2015). limma powers differential expression analyses for RNA-sequencing and microarray studies. Nucleic Acids Research, 43, e47–e47.

Rizzini L., Favory J.-J., Cloix C., Faggionato D., O’Hara A., Kaiserli E., Baumeister R., Schäfer E., Nagy F., Jenkins G.I., et al. (2011). Perception of UV-B by the Arabidopsis UVR8 protein. Science, 332, 103–106.

Robinson M.D., McCarthy D.J., Smyth G.K. (2010). edgeR: a Bioconductor package for differential expression analysis of digital gene expression data. Bioinformatics, 26, 139–140.

Sainsbury F., Thuenemann E.C., Lomonossoff G.P. (2009). pEAQ: versatile expression vectors for easy and quick transient expression of heterologous proteins in plants. Plant Biotechnology Journal, 7, 682–693.

Sakai R., Winand R., Verbeiren T., Moere A.V., Aerts J. (2014). dendsort: modular leaf ordering methods for dendrogram representations in R. F1000Research 3: 177.

Siipola S.M., Kotilainen T., Sipari N., Morales L.O., Lindfors A.V., Robson T.M., Aphalo P.J. (2015). Epidermal UV-A absorbance and whole-leaf flavonoid composition in pea respond more to solar blue light than to solar UV radiation. Plant, Cell and Environment, 38, 941–952.

Stracke R., Ishihara H., Huep G., Barsch A., Mehrtens F., Niehaus K., Weisshaar B. (2007). Differential regulation of closely related R2R3-MYB transcription factors controls flavonol accumulation in different parts of the *Arabidopsis thaliana* seedling. The Plant Journal, 50, 660–677.

Wang W., Lu X., Li L., Lian H., Mao Z., Xu P., Guo T., Xu F., Du S., Cao X., et al. (2018). Photoexcited CRYPTOCHROME1 interacts with dephosphorylated BES1 to regulate brassinosteroid signaling and photomorphogenesis in Arabidopsis. The Plant Cell, 30, 1989–2005.

Wang H., Ma L.G., Li J.M., Zhao H.Y., Deng X.W. (2001). Direct interaction of Arabidopsis Cryptochromes with COP1 in light control development. Science, 294, 154–158.

Wang Q., Zuo Z., Wang X., Gu L., Yoshizumi T., Yang Z., Yang L., Liu Q., Liu W., Han Y.-J., et al. (2016). Photoactivation and inactivation of *Arabidopsis* cryptochrome 2. Science, 354, 343–347.

Warnes G.R., Bolker B., Lumley T., Johnson R.C. (2018). gmodels: Various R programming tools for model fitting version 2.18.1 from CRAN. URL https://cran.r-project.org/web/packages/gmodels/index.html [accessed 12 June 2019].

Wickham H. (2009). Ggplot2c: elegant graphics for data analysis. Springer.

Wu D., Hu Q., Yan Z., Chen W., Yan C., Huang X., Zhang J., Yang P., Deng H., Wang J., et al. (2012). Structural basis of ultraviolet-B perception by UVR8. Nature, 484, 214–219.

Yang Y., Liang T., Zhang L., Shao K., Gu X., Shang R., Shi N., Li X., Zhang P., Liu H. (2018). UVR8 interacts with WRKY36 to regulate HY5 transcription and hypocotyl elongation in Arabidopsis. Nature Plants, 4, 98–107.

Yang Z., Liu B., Su J., Liao J., Lin C., Oka Y. (2017). Cryptochromes orchestrate transcription regulation of diverse blue light responses in plants. Photochemistry and Photobiology, 93, 112–127.

Yang X., Montano S., Ren Z. (2015). How does photoreceptor UVR8 perceive a UV-B Signal? Photochemistry and Photobiology, 91, 993–1003.

Yang H.-Q., Tang R.-H., Cashmore A.R. (2001). The signaling mechanism of Arabidopsis CRY1 involves direct interaction with COP1. The Plant Cell, 13, 2573–2587.

Yu X., Liu H., Klejnot J., Lin C. (2010). The Cryptochrome blue light receptors. The Arabidopsis Book, 8, e0135.

Zhang R., Calixto C.P.G., Marquez Y., Venhuizen P., Tzioutziou N.A., Guo W., Spensley M., Entizne J.C., Lewandowska D., Ten Have S., et al. (2017). A high quality Arabidopsis transcriptome for accurate transcript-level analysis of alternative splicing. Nucleic Acids Research, 45, 5061–5073.

Zuo Z., Liu H., Liu B., Liu X., Lin C. (2011). Blue light-dependent interaction of CRY2 with SPA1 regulates COP1 activity and floral initiation in Arabidopsis. Current Biology, 21, 841–847.

