## Supplemental Information for "The photoreceptor UVR8 mediates the perception of both UV-B and UV-A wavelengths up to 350 nm of sunlight with responsivity moderated by cryptochromes"

**The following Supporting Information is available for this article:**

Figure S1. Solar spectrum at different times of the day when plants were moved outdoors.

Figure S2. Photon irradiance for different wavebands in solar radiation.

Figure S3. Multidimensional scaling of RNA-seq data.

Figure S4. Comparison between RNA-seq and qRT-PCR data.

Figure S5. Venn diagrams showing the number differentially expressed genes in RNA-seq data.

Figure S6. Enrichment of KEGG pathways in RNA-seq data.

Figure S7. *In vitro* absorption spectra of Arabidopsis UVR8 protein.

Figure S8. Position weight matrices of the enriched DNA-binding motifs.

Figure S9. Transcript abundance of seven genes measured using qRT-PCR.

Table S1. Information of primers used and genes assessed in qRT-PCR.

Table S2. Summary of the ANOVA of the qRT-PCR data.

Methods S1. Description of the filters and the waveband contrasts.

Dataset S1. Outcome of differential gene expression analysis for the three genotypes and multiple waveband contrasts combination. The dataset is included as a separate file in .Rda format and can be read using R.

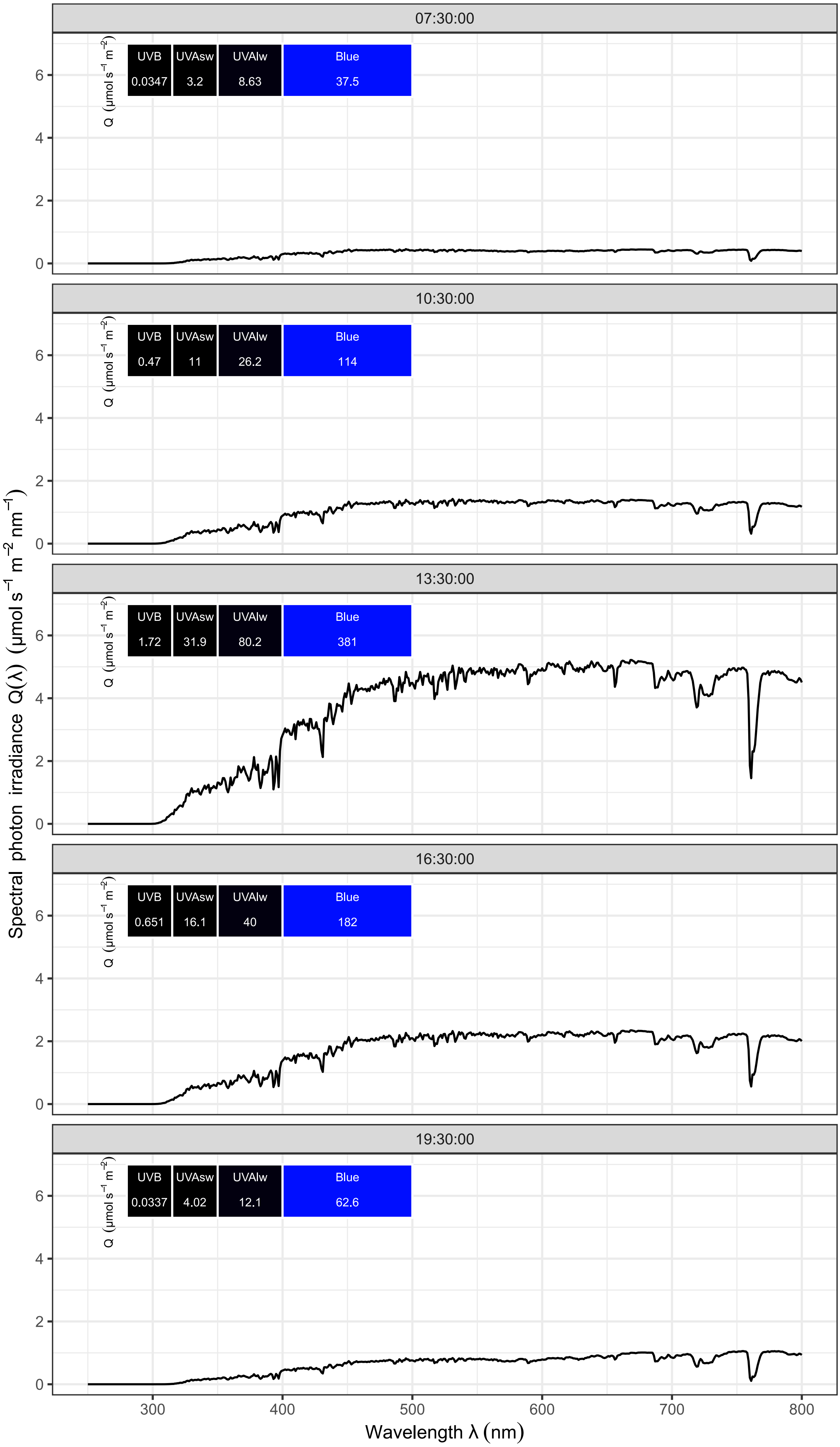

Fig. S1. Solar spectrum at different times of the day on 21 August 2014, the day when plants were moved outdoors for filter treatments. The labels show the photons irradiance of waveband contrasts used in the experiment.

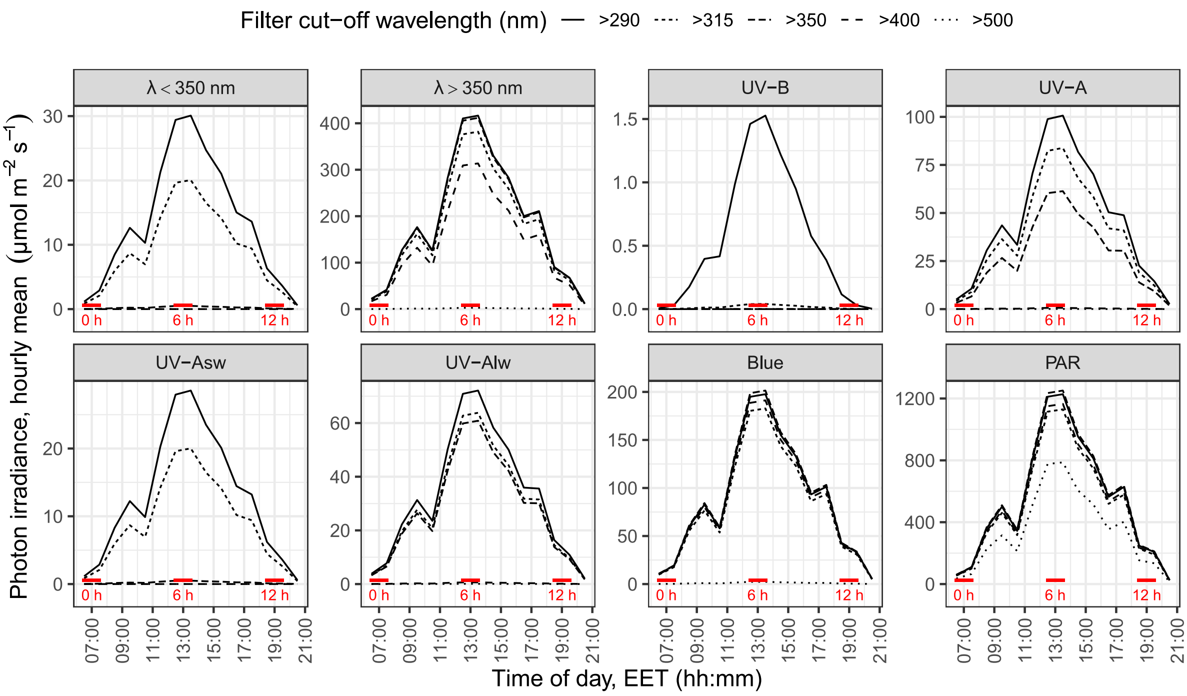

Fig. S2. Photon irradiance of solar radiation for the wavebands λ < 350 nm (290–350 nm), λ > 350 nm (350–500 nm), UV-B (290–315 nm), UV-A (315–400 nm), UV-$A_{sw}$ (315–350 nm), UV-$A_{lw}$ (350–400 nm), blue (400–500 nm) and PAR (400–700 nm) throughout the day. Values were estimated for each filter treatment as hourly means for the period of plants exposure to sunlight. The red horizontal lines show when the plants were moved outdoors (0 h) or sampled (6 h and 12 h).

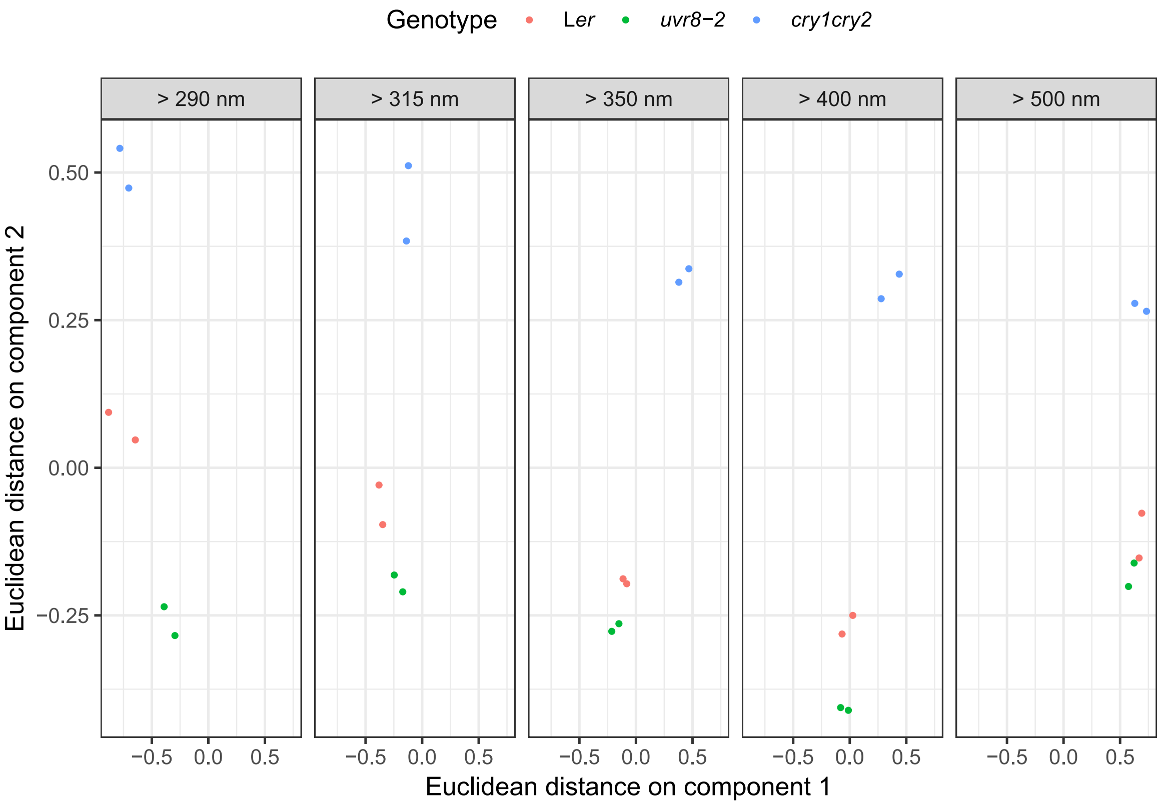

Fig. S3. Multidimensional scaling (MDS) plot showing variation among RNA-seq libraries used in the experiment. Two libraries per treatment per genotype were constructed by pooling two pairs of RNA samples extracted from four biological replicates. The first MDS component separated the filter treatments and the second component the genotypes.

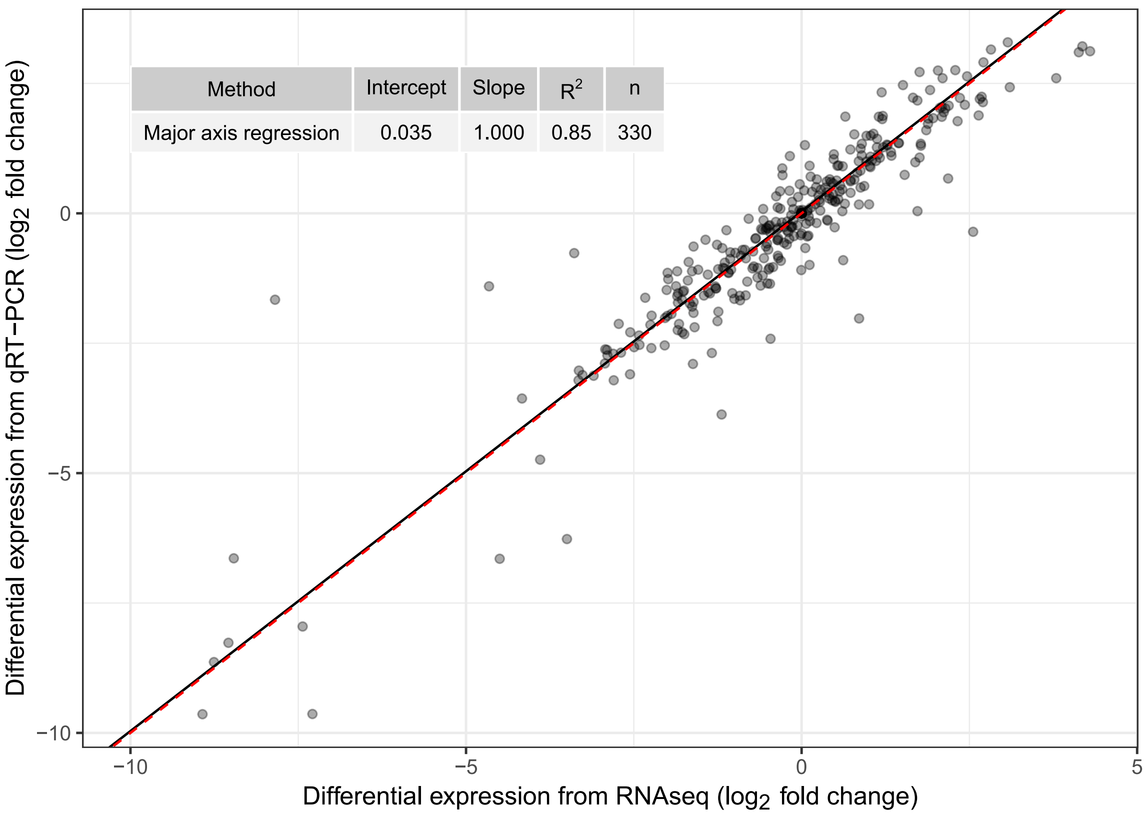

Fig. S4. Comparison between the two methods used to assess the differential gene expression after 6 h of solar treatments. For comparison $\log_2$FC was used for both data sets and rescaled using the same normalization. The whole line shows the fitted main axis linear regression, the red dashed line shows 1:1 slope and the inset the estimates for the parameters.

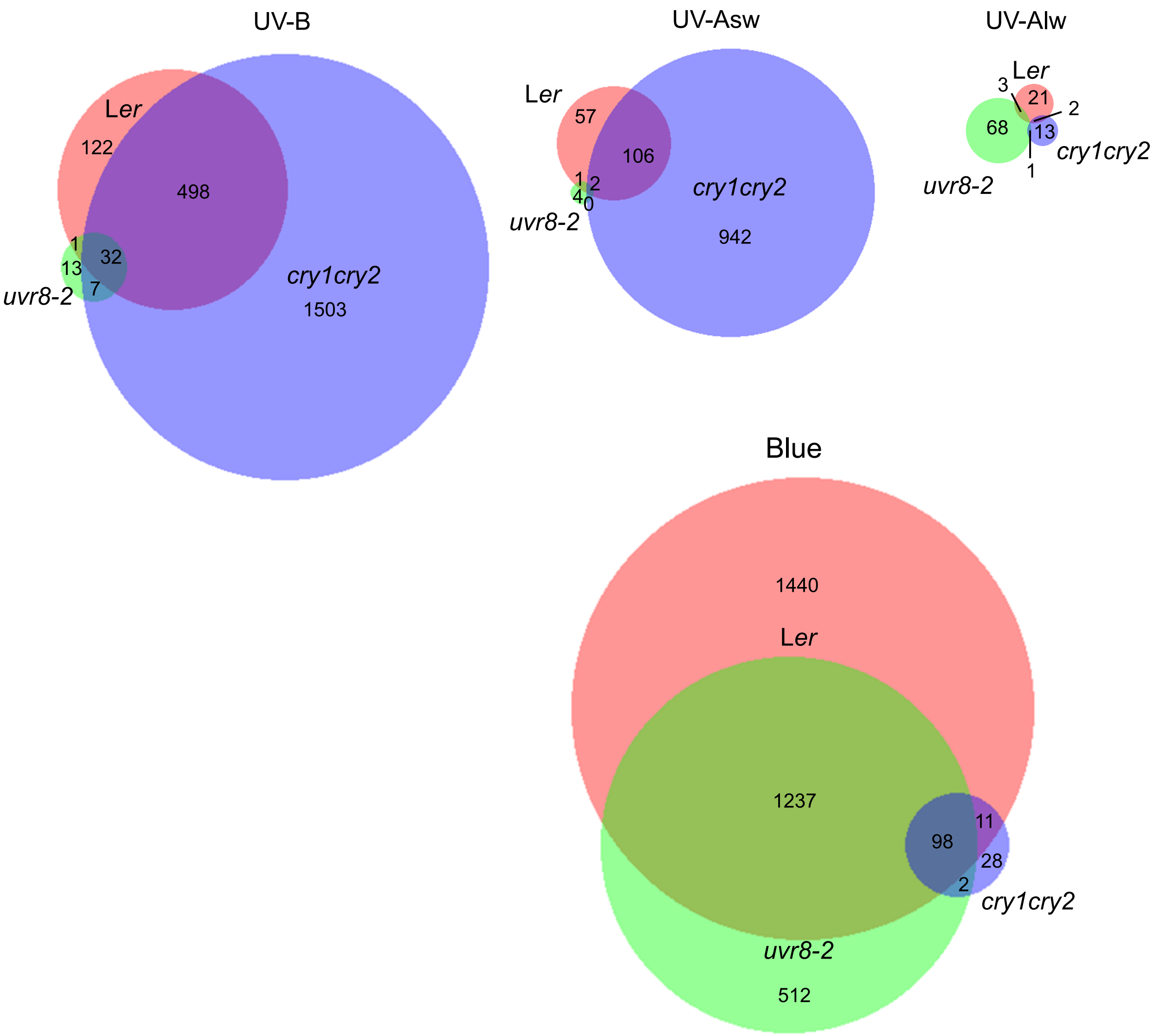

Fig. S5. Venn diagrams showing number of DEGs in L*er*, *uvr8-2* and *cry1cry2* in response to 6 h of solar UV-B, UV-A$_{sw}$, UV-A$_{lw}$ and blue. FC > 1.5, $P_{adjust} < 0.05$.

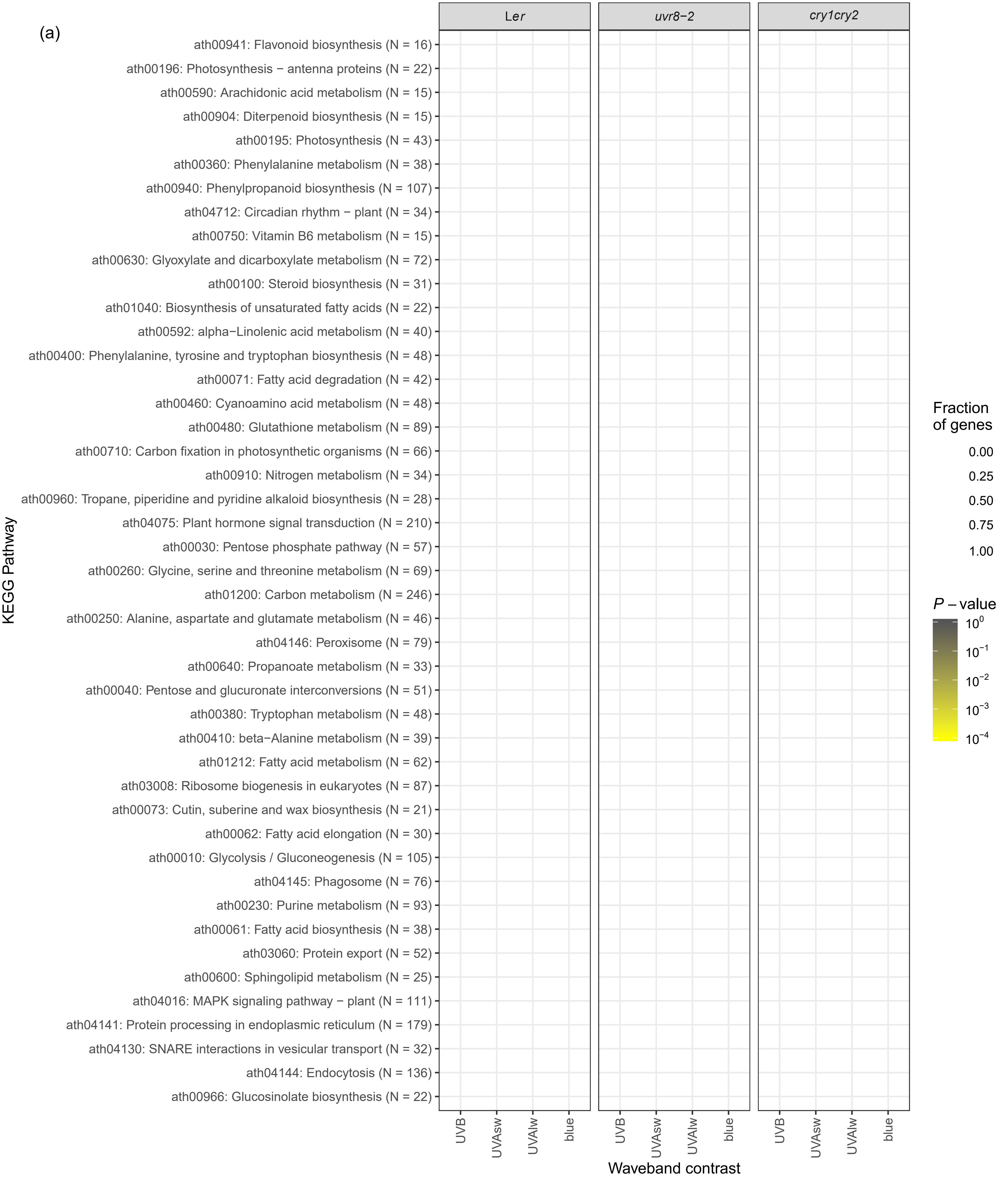
(a)
Ler
uvr8−2
cry1cry2
ath00941: Flavonoid biosynthesis (N = 16)
ath00196: Photosynthesis − antenna proteins (N = 22)
ath00590: Arachidonic acid metabolism (N = 15)
ath00904: Diterpenoid biosynthesis (N = 15)
ath00195: Photosynthesis (N = 43)
ath00360: Phenylalanine metabolism (N = 38)
ath00940: Phenylpropanoid biosynthesis (N = 107)
ath04712: Circadian rhythm − plant (N = 34)
ath00750: Vitamin B6 metabolism (N = 15)
ath00630: Glyoxylate and dicarboxylate metabolism (N = 72)
ath00100: Steroid biosynthesis (N = 31)
ath01040: Biosynthesis of unsaturated fatty acids (N = 22)
ath00592: alpha−Linolenic acid metabolism (N = 40)
ath00400: Phenylalanine, tyrosine and tryptophan biosynthesis (N = 48)
ath00071: Fatty acid degradation (N = 42)
ath00460: Cyanoamino acid metabolism (N = 48)
ath00480: Glutathione metabolism (N = 89)
ath00710: Carbon fixation in photosynthetic organisms (N = 66)
ath00910: Nitrogen metabolism (N = 34)
ath00960: Tropane, piperidine and pyridine alkaloid biosynthesis (N = 28)
ath04075: Plant hormone signal transduction (N = 210)
ath00030: Pentose phosphate pathway (N = 57)
ath00260: Glycine, serine and threonine metabolism (N = 69)
ath01200: Carbon metabolism (N = 246)
ath00250: Alanine, aspartate and glutamate metabolism (N = 46)
ath04146: Peroxisome (N = 79)
ath00640: Propanoate metabolism (N = 33)
ath00040: Pentose and glucuronate interconversions (N = 51)
ath00380: Tryptophan metabolism (N = 48)
ath00410: beta−Alanine metabolism (N = 39)
ath01212: Fatty acid metabolism (N = 62)
ath03008: Ribosome biogenesis in eukaryotes (N = 87)
ath00073: Cutin, suberine and wax biosynthesis (N = 21)
ath00062: Fatty acid elongation (N = 30)
ath00010: Glycolysis / Gluconeogenesis (N = 105)
ath04145: Phagosome (N = 76)
ath00230: Purine metabolism (N = 93)
ath00061: Fatty acid biosynthesis (N = 38)
ath03060: Protein export (N = 52)
ath00600: Sphingolipid metabolism (N = 25)
ath04016: MAPK signaling pathway − plant (N = 111)
ath04141: Protein processing in endoplasmic reticulum (N = 179)
ath04130: SNARE interactions in vesicular transport (N = 32)
ath04144: Endocytosis (N = 136)
ath00966: Glucosinolate biosynthesis (N = 22)
KEGG Pathway
UVB
UVAsw
UVAlw
blue
Waveband contrast
Fraction
of genes
0.00
0.25
0.50
0.75
1.00
P − value
10^0
10^−1
10^−2
10^−3
10^−4

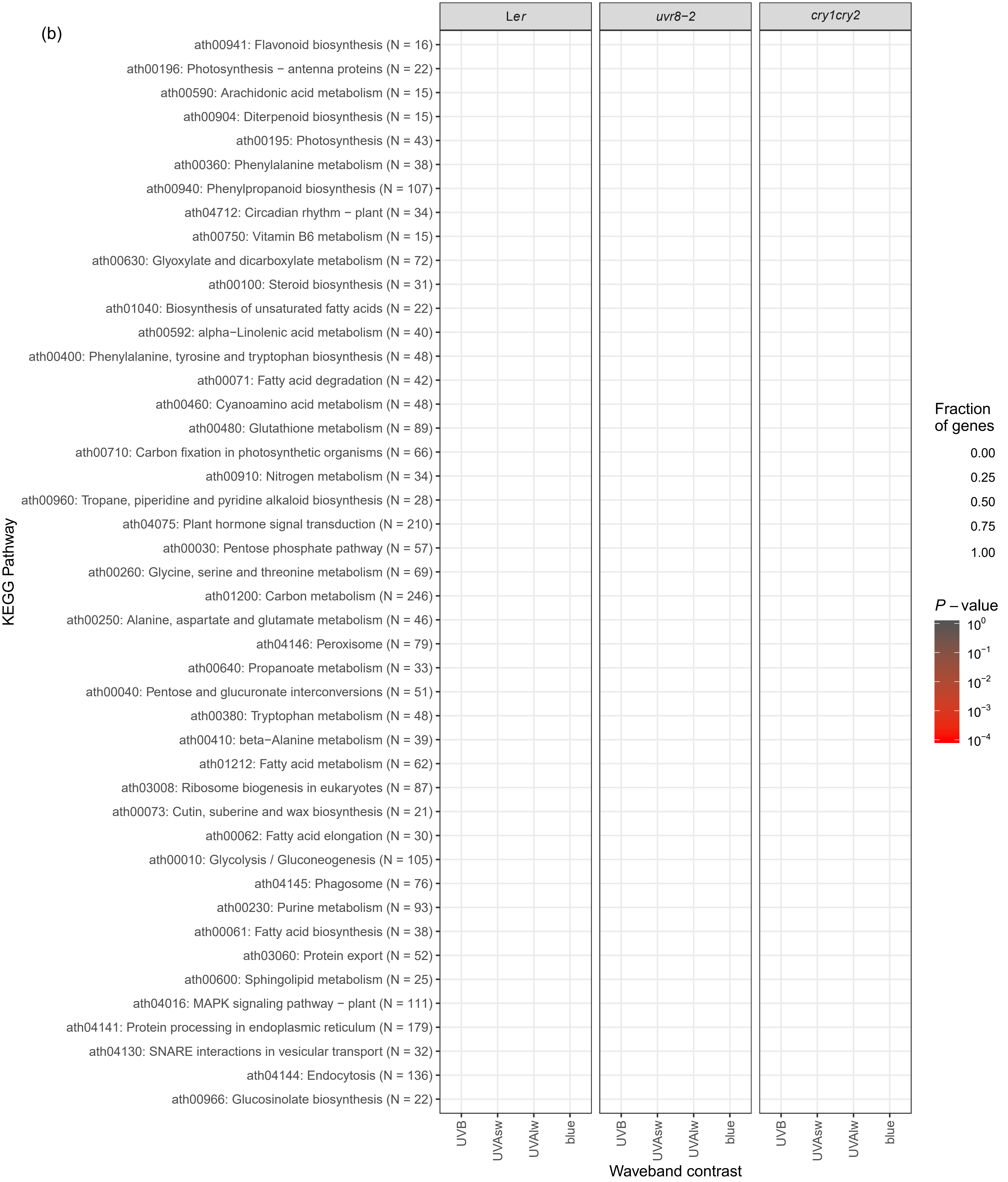
(b)
Ler
uvr8−2
cry1cry2
ath00941: Flavonoid biosynthesis (N = 16)
ath00196: Photosynthesis − antenna proteins (N = 22)
ath00590: Arachidonic acid metabolism (N = 15)
ath00904: Diterpenoid biosynthesis (N = 15)
ath00195: Photosynthesis (N = 43)
ath00360: Phenylalanine metabolism (N = 38)
ath00940: Phenylpropanoid biosynthesis (N = 107)
ath04712: Circadian rhythm − plant (N = 34)
ath00750: Vitamin B6 metabolism (N = 15)
ath00630: Glyoxylate and dicarboxylate metabolism (N = 72)
ath00100: Steroid biosynthesis (N = 31)
ath01040: Biosynthesis of unsaturated fatty acids (N = 22)
ath00592: alpha−Linolenic acid metabolism (N = 40)
ath00400: Phenylalanine, tyrosine and tryptophan biosynthesis (N = 48)
ath00071: Fatty acid degradation (N = 42)
ath00460: Cyanoamino acid metabolism (N = 48)
ath00480: Glutathione metabolism (N = 89)
ath00710: Carbon fixation in photosynthetic organisms (N = 66)
ath00910: Nitrogen metabolism (N = 34)
ath00960: Tropane, piperidine and pyridine alkaloid biosynthesis (N = 28)
ath04075: Plant hormone signal transduction (N = 210)
ath00030: Pentose phosphate pathway (N = 57)
ath00260: Glycine, serine and threonine metabolism (N = 69)
ath01200: Carbon metabolism (N = 246)
ath00250: Alanine, aspartate and glutamate metabolism (N = 46)
ath04146: Peroxisome (N = 79)
ath00640: Propanoate metabolism (N = 33)
ath00040: Pentose and glucuronate interconversions (N = 51)
ath00380: Tryptophan metabolism (N = 48)
ath00410: beta−Alanine metabolism (N = 39)
ath01212: Fatty acid metabolism (N = 62)
ath03008: Ribosome biogenesis in eukaryotes (N = 87)
ath00073: Cutin, suberine and wax biosynthesis (N = 21)
ath00062: Fatty acid elongation (N = 30)
ath00010: Glycolysis / Gluconeogenesis (N = 105)
ath04145: Phagosome (N = 76)
ath00230: Purine metabolism (N = 93)
ath00061: Fatty acid biosynthesis (N = 38)
ath03060: Protein export (N = 52)
ath00600: Sphingolipid metabolism (N = 25)
ath04016: MAPK signaling pathway − plant (N = 111)
ath04141: Protein processing in endoplasmic reticulum (N = 179)
ath04130: SNARE interactions in vesicular transport (N = 32)
ath04144: Endocytosis (N = 136)
ath00966: Glucosinolate biosynthesis (N = 22)
KEGG Pathway
UVB
UVAsw
UVAlw
blue
Waveband contrast
Fraction of genes
0.00
0.25
0.50
0.75
1.00
P − value
10^0
10^−1
10^−2
10^−3
10^−4

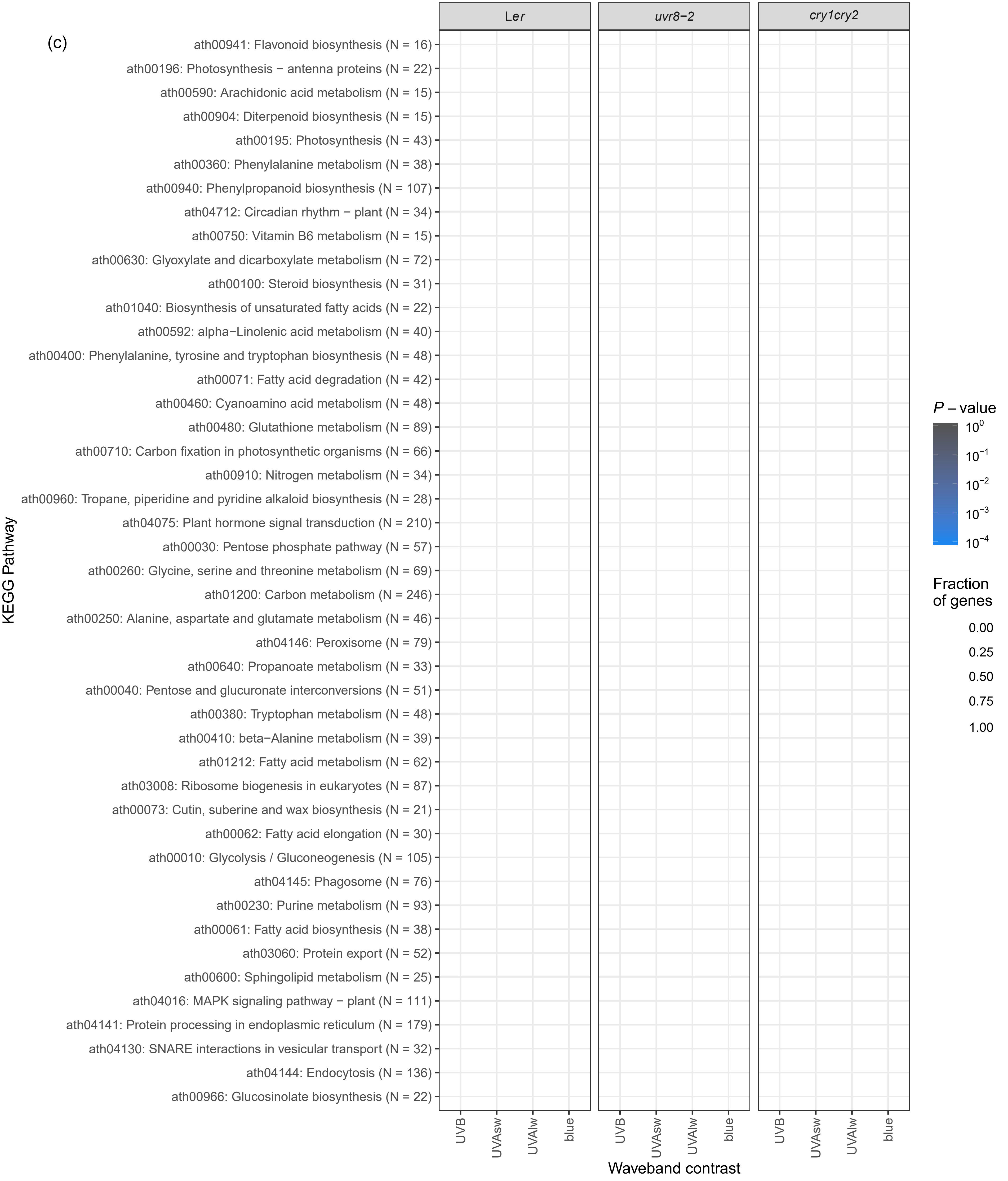

(c)
Ler
uvr8−2
cry1cry2
ath00941: Flavonoid biosynthesis (N = 16)
ath00196: Photosynthesis − antenna proteins (N = 22)
ath00590: Arachidonic acid metabolism (N = 15)
ath00904: Diterpenoid biosynthesis (N = 15)
ath00195: Photosynthesis (N = 43)
ath00360: Phenylalanine metabolism (N = 38)
ath00940: Phenylpropanoid biosynthesis (N = 107)
ath04712: Circadian rhythm − plant (N = 34)
ath00750: Vitamin B6 metabolism (N = 15)
ath00630: Glyoxylate and dicarboxylate metabolism (N = 72)
ath00100: Steroid biosynthesis (N = 31)
ath01040: Biosynthesis of unsaturated fatty acids (N = 22)
ath00592: alpha−Linolenic acid metabolism (N = 40)
ath00400: Phenylalanine, tyrosine and tryptophan biosynthesis (N = 48)
ath00071: Fatty acid degradation (N = 42)
ath00460: Cyanoamino acid metabolism (N = 48)
ath00480: Glutathione metabolism (N = 89)
ath00710: Carbon fixation in photosynthetic organisms (N = 66)
ath00910: Nitrogen metabolism (N = 34)
ath00960: Tropane, piperidine and pyridine alkaloid biosynthesis (N = 28)
ath04075: Plant hormone signal transduction (N = 210)
ath00030: Pentose phosphate pathway (N = 57)
ath00260: Glycine, serine and threonine metabolism (N = 69)
ath01200: Carbon metabolism (N = 246)
ath00250: Alanine, aspartate and glutamate metabolism (N = 46)
ath04146: Peroxisome (N = 79)
ath00640: Propanoate metabolism (N = 33)
ath00040: Pentose and glucuronate interconversions (N = 51)
ath00380: Tryptophan metabolism (N = 48)
ath00410: beta−Alanine metabolism (N = 39)
ath01212: Fatty acid metabolism (N = 62)
ath03008: Ribosome biogenesis in eukaryotes (N = 87)
ath00073: Cutin, suberine and wax biosynthesis (N = 21)
ath00062: Fatty acid elongation (N = 30)
ath00010: Glycolysis / Gluconeogenesis (N = 105)
ath04145: Phagosome (N = 76)
ath00230: Purine metabolism (N = 93)
ath00061: Fatty acid biosynthesis (N = 38)
ath03060: Protein export (N = 52)
ath00600: Sphingolipid metabolism (N = 25)
ath04016: MAPK signaling pathway − plant (N = 111)
ath04141: Protein processing in endoplasmic reticulum (N = 179)
ath04130: SNARE interactions in vesicular transport (N = 32)
ath04144: Endocytosis (N = 136)
ath00966: Glucosinolate biosynthesis (N = 22)
KEGG Pathway
UVB
UVAsw
UVAlw
blue
Waveband contrast
P − value
10^0
10^−1
10^−2
10^−3
10^−4
Fraction of genes
0.00
0.25
0.50
0.75
1.00

Fig. S6. KEGG pathway analysis. Enriched pathways for (a) all the DEGs (yellow circles), (b) DEGs with increased expression (red circles), (c) DEGs with decreased expression (blue circles). KEGG pathways with *P*-value $< 0.01$ for at least one contrast and genotype combination are included in the figure. The area of the circles shows a fraction of DEGs in a particular KEGG pathway against the total number of genes annotated for that KEGG pathway. The saturation of the color of circles shows *P*-value on a log scale after setting all the *P*-values $<10^{-4}$ to $10^{-4}$. N refers to the total number of genes annotated for that KEGG pathway in the database. The pathways were ordered by the average of fractions of DEGs, therefore, the most responsive pathways to waveband contrasts come at the top of the figure.

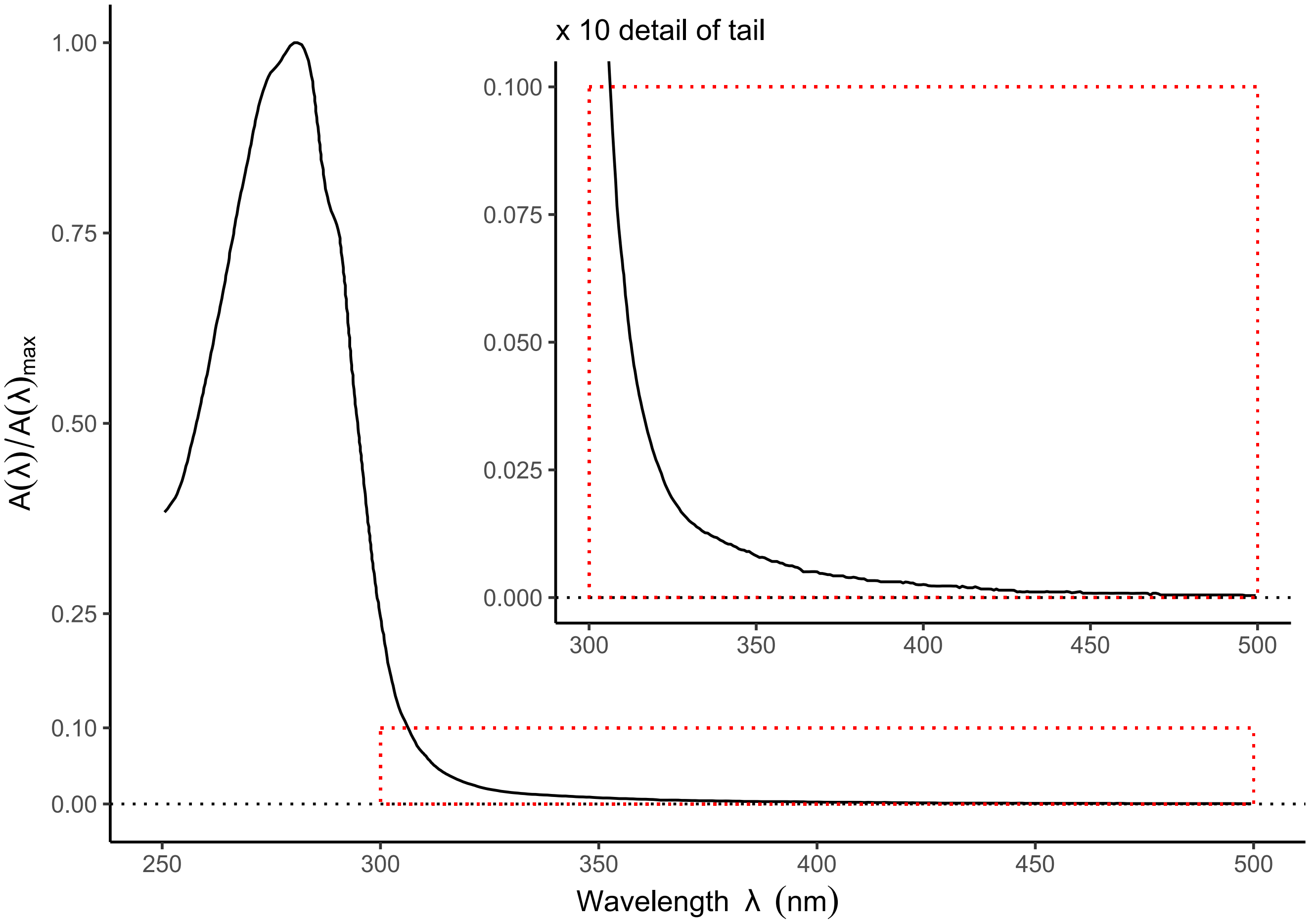

Fig. S7. *In vitro* absorption spectrum of *Arabidopsis thaliana* UVR8 protein expressed and purified from *Escherichia coli.* To improve the signal to noise ratio for spectral regions covering both long and short wavelengths, two separate datasets were recorded for both high and medium concentrated UVR8 protein samples at 57.0 µM and 4.5 µM, respectively. The spectrum shown is the result of splicing these separately measured spectra. After splicing the spectrum was normalized for expression in relative spectral absorbance units.

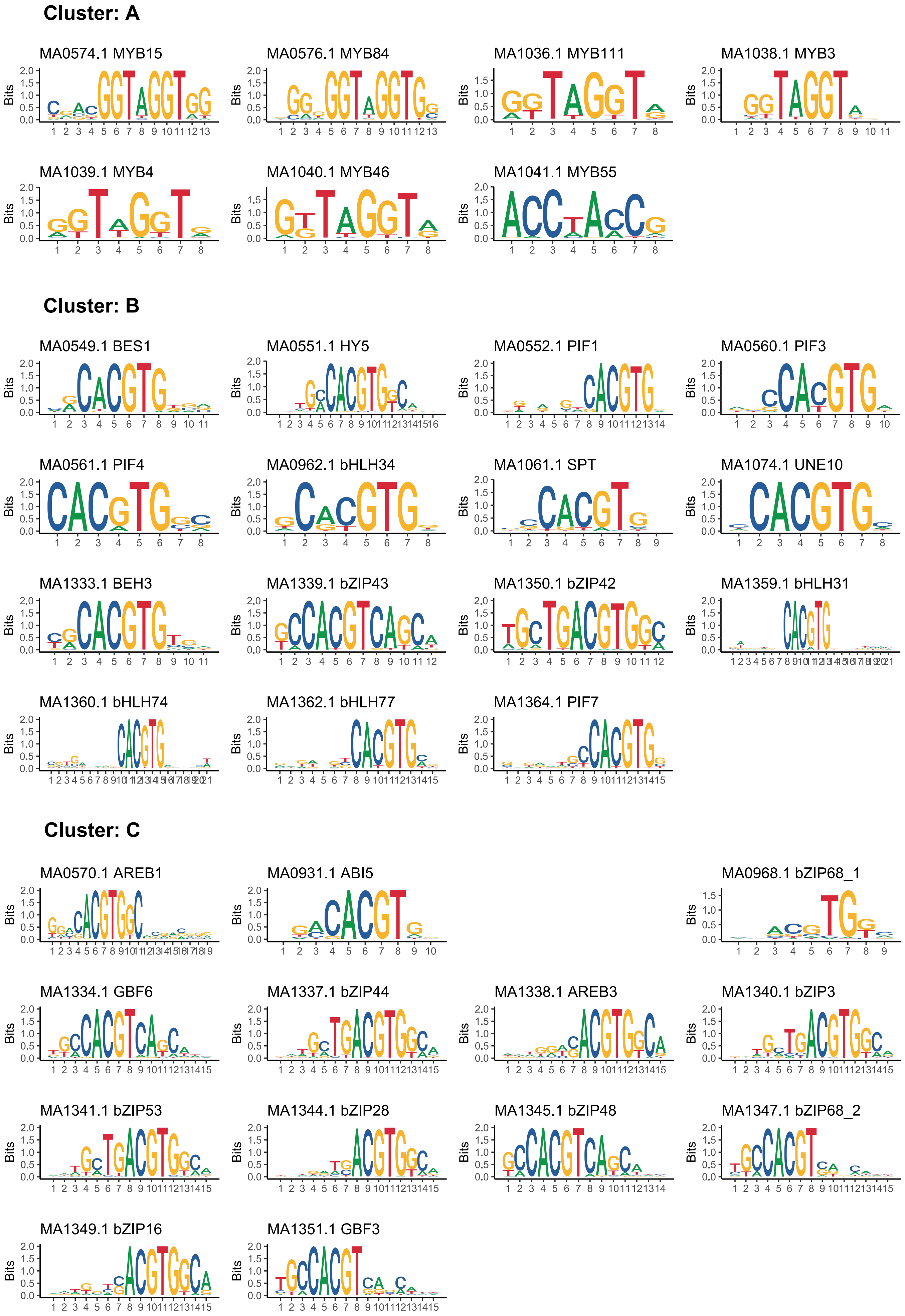
Cluster: A
MA0574.1 MYB15
MA0576.1 MYB84
MA1036.1 MYB111
MA1038.1 MYB3
MA1039.1 MYB4
MA1040.1 MYB46
MA1041.1 MYB55
Cluster: B
MA0549.1 BES1
MA0551.1 HY5
MA0552.1 PIF1
MA0560.1 PIF3
MA0561.1 PIF4
MA0962.1 bHLH34
MA1061.1 SPT
MA1074.1 UNE10
MA1333.1 BEH3
MA1339.1 bZIP43
MA1350.1 bZIP42
MA1359.1 bHLH31
MA1360.1 bHLH74
MA1362.1 bHLH77
MA1364.1 PIF7
Cluster: C
MA0570.1 AREB1
MA0931.1 ABI5
MA0968.1 bZIP68_1
MA1334.1 GBF6
MA1337.1 bZIP44
MA1338.1 AREB3
MA1340.1 bZIP3
MA1341.1 bZIP53
MA1344.1 bZIP28
MA1345.1 bZIP48
MA1347.1 bZIP68_2
MA1349.1 bZIP16
MA1351.1 GBF3
Bits

### Cluster: D

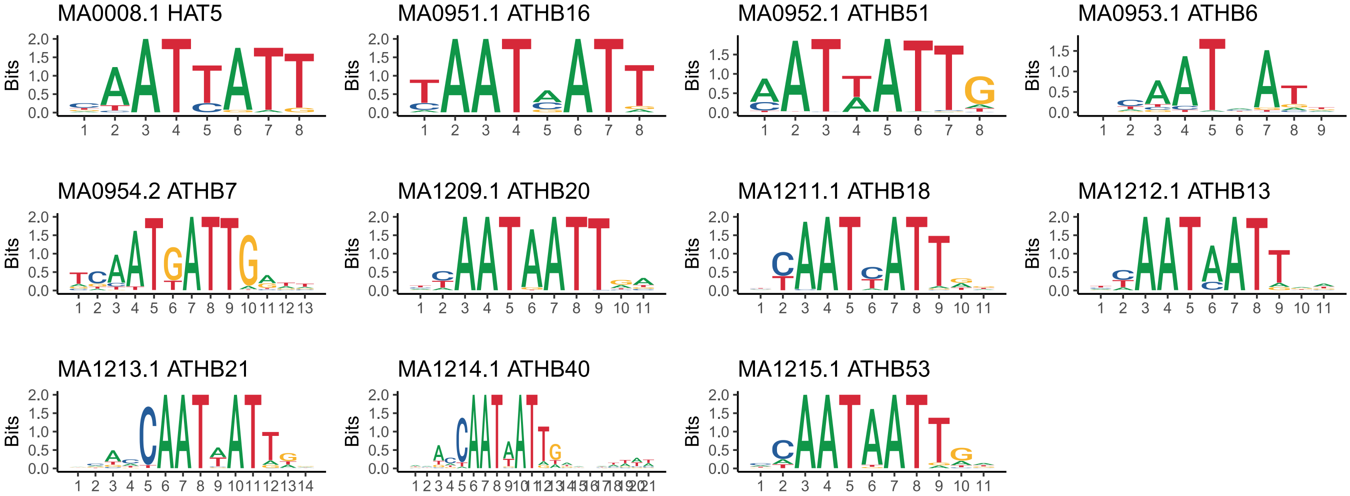

### Cluster: E

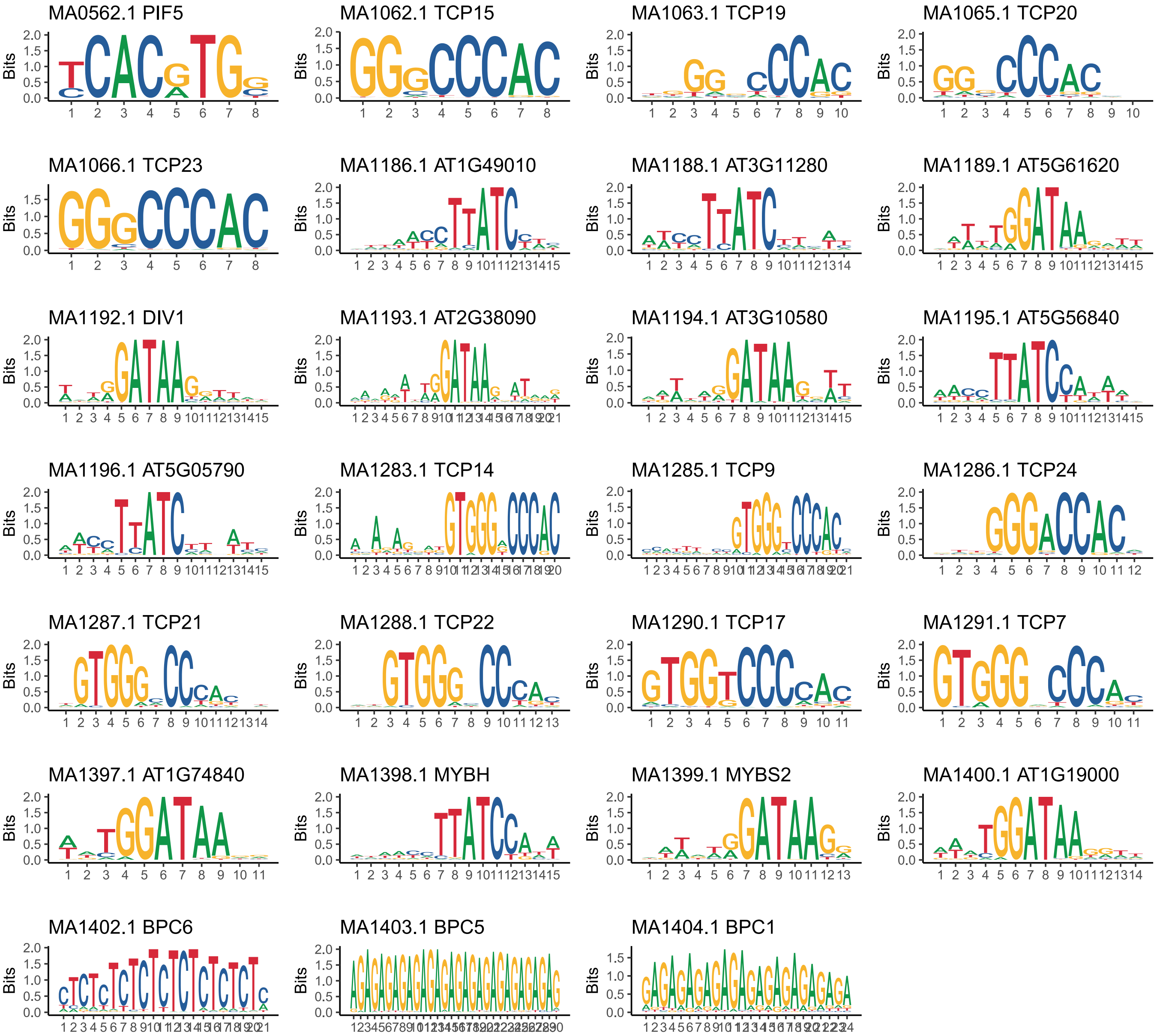

### Cluster: F

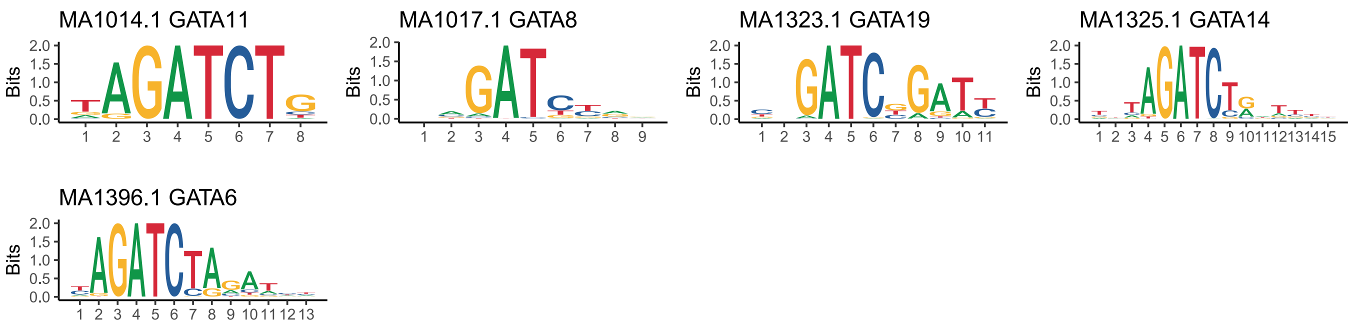

### Cluster: G

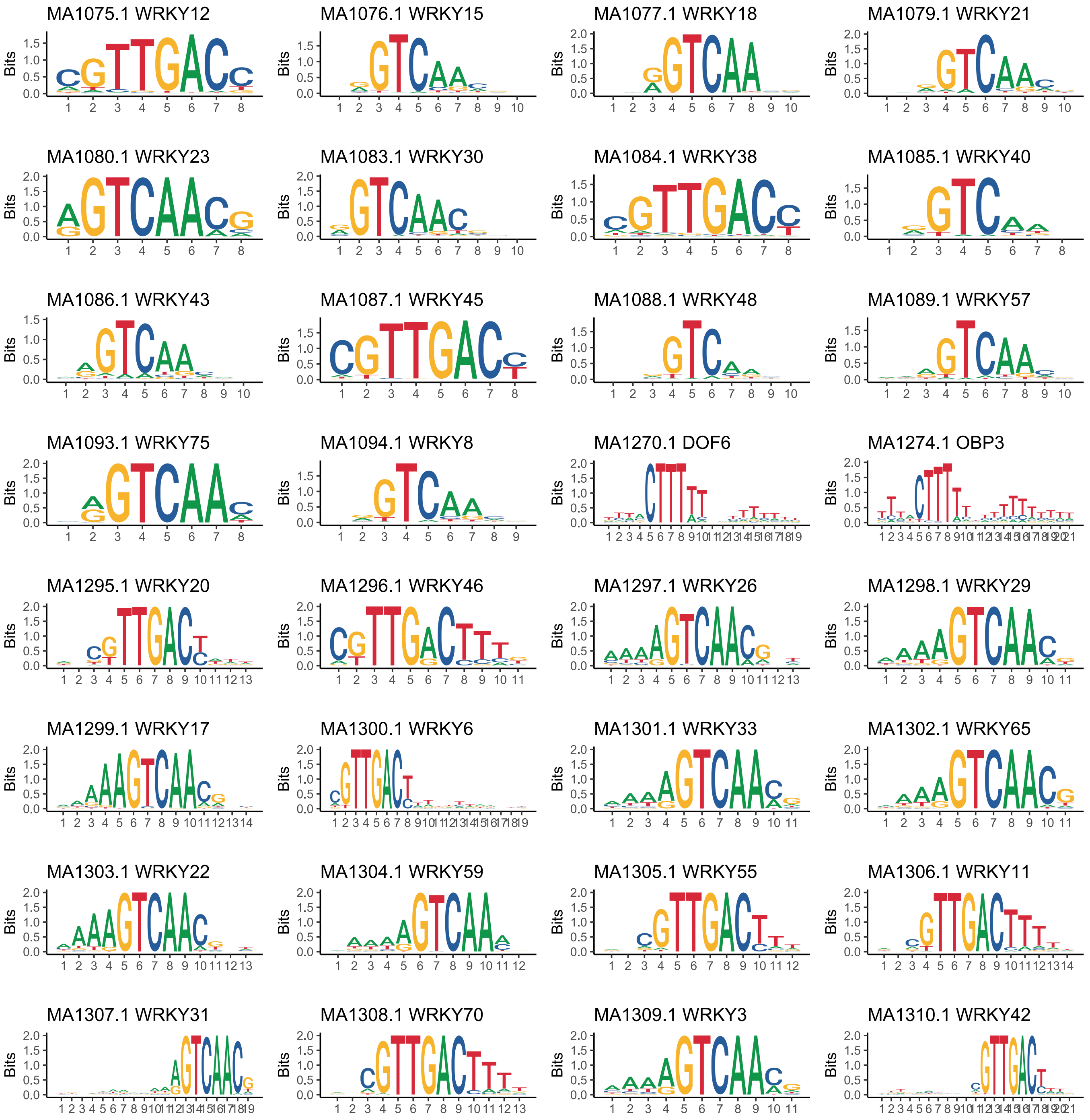

Cluster G continues on next page.

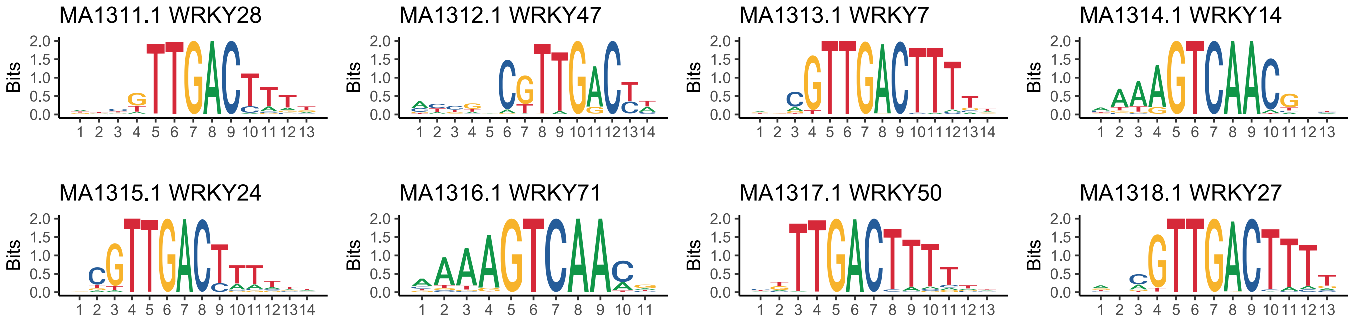

**Cluster: H**

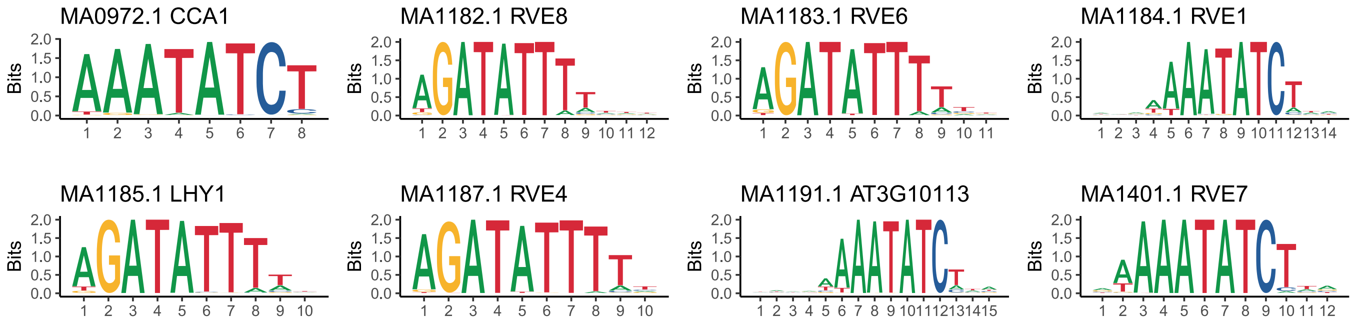

**Cluster: I**

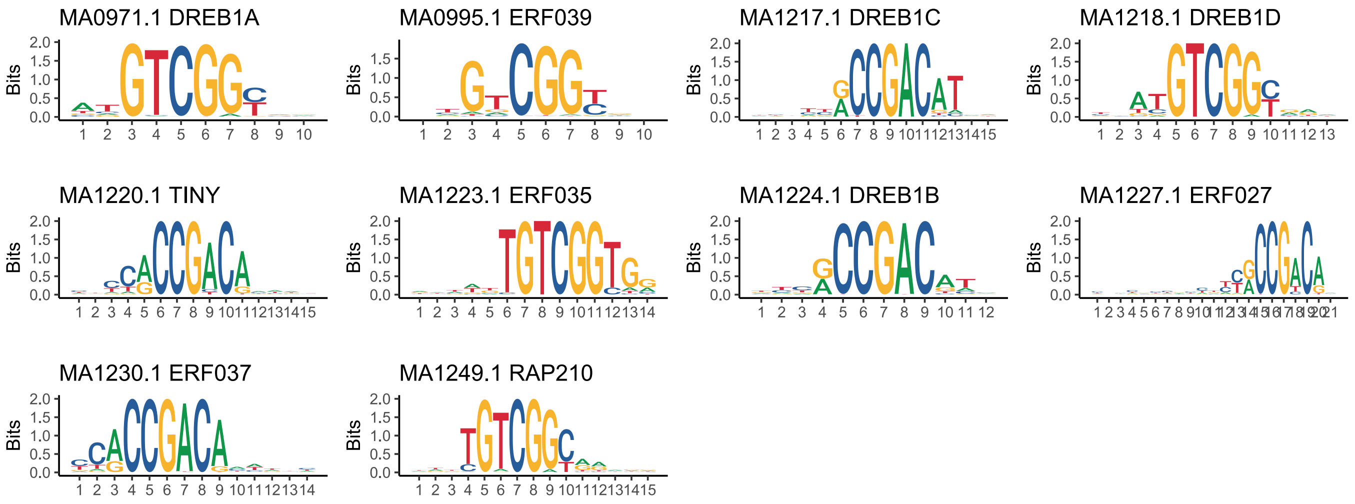

**Cluster: J**

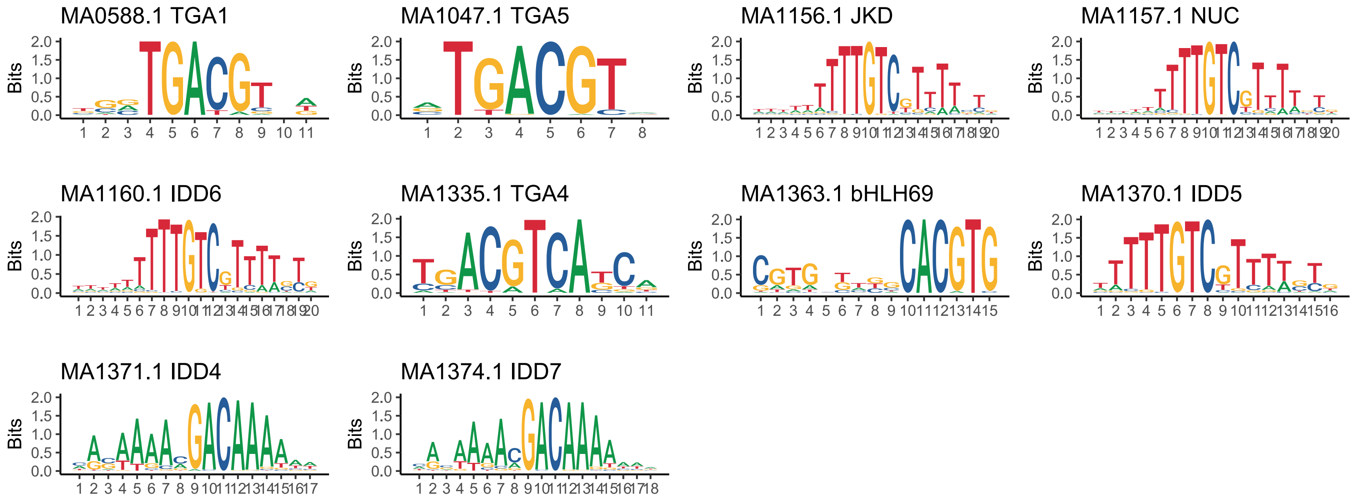

### Cluster: K

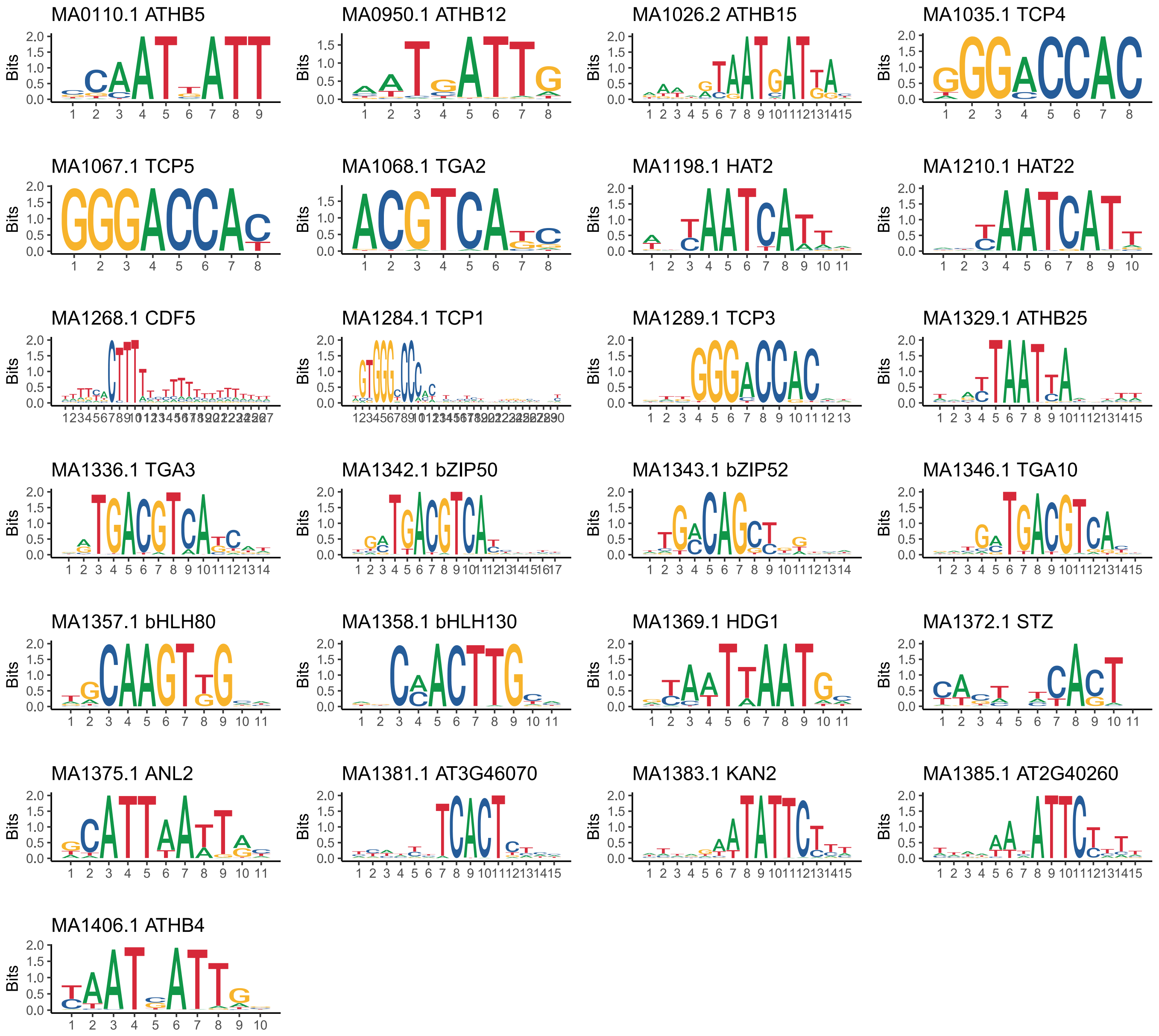

### Cluster: L

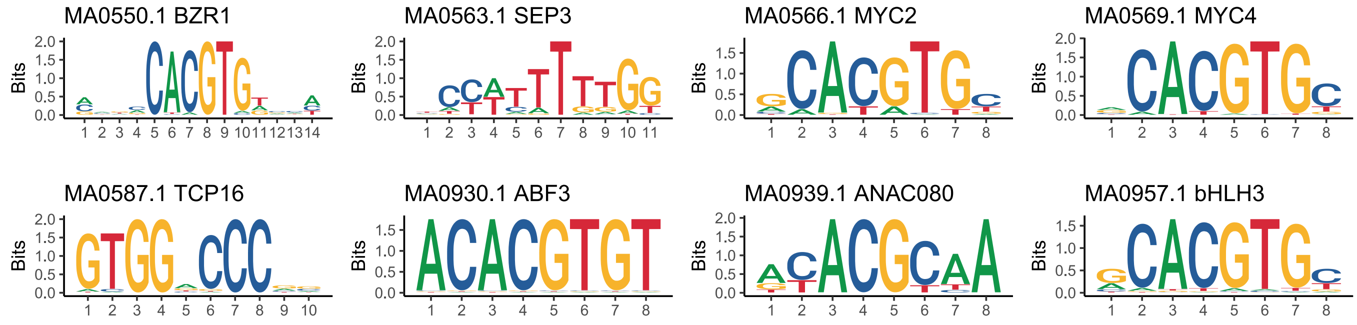

Cluster L continues on next page.

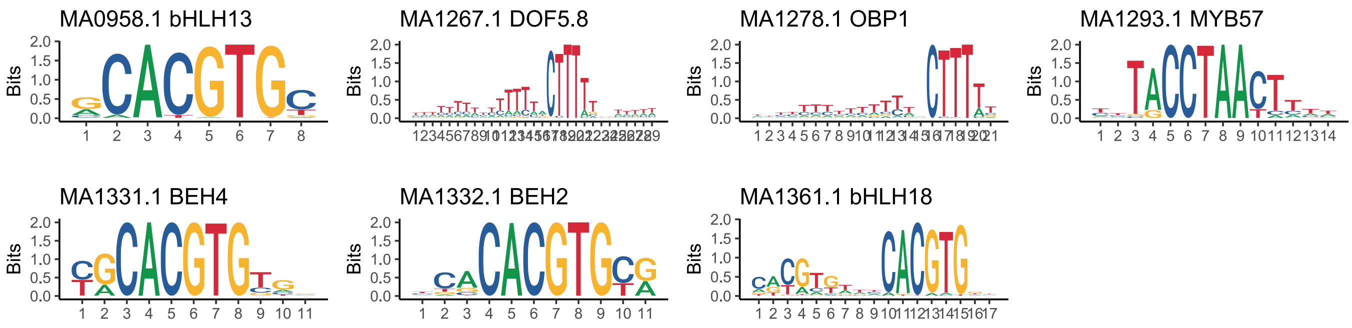

Fig. S8. Position weight matrices of the enriched DNA-binding motifs. The motifs were obtained from JASPAR 2018 database. JASPAR ids and gene symbols are shown in the figure.

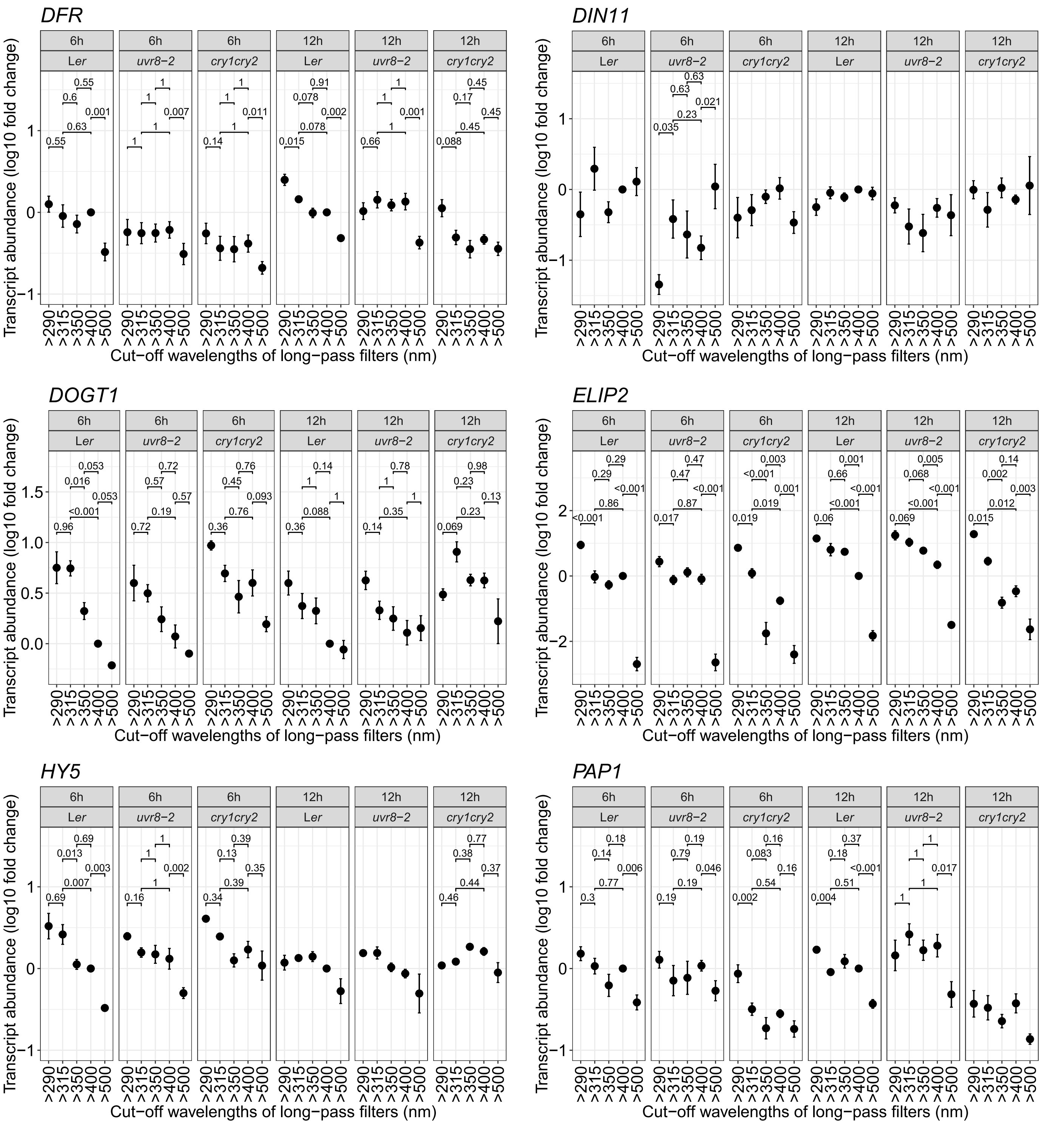

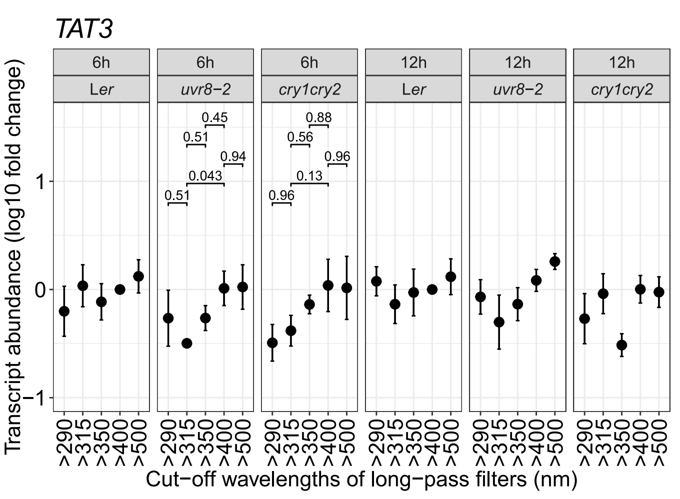

Fig. S9. Transcript abundance of seven marker genes measured with qRT-PCR after 6 h and 12 h of solar treatments. Mean1±SE from four biological replicates. The horizontal bars show $P_{adjust}$ values for pair-wise comparisons between treatments within each genotype. $P_{adjust}$ values are shown in those panels where the overall effect of filter treatments within a genotype and time point was significant (see Table S2).

Table S1. Information of primers used and genes assessed in qRT-PCR.

| **Gene** | **Annotation** | **Forward primer** | **Reverse primer** | **Amplification efficiency** |
|---|---|---|---|---|
| *CHALCONE ISOMERASE* (*CHI*) | AT3G55120 | TCCCTGAAACCGGGATCGCTGT | AGTGCCAGGTGACACACCGT | 2 |
| *CHALCONE SYNTHASE* (*CHS*) | AT5G13930 | ATGTCGAGCGCGTGCGTTCT | TCTCCTGTCGTGGCCACACCA | 2 |
| *DIHYDROFLAVONOL REDUCTASE* (*DFR*) | AT5G42800 | ACCGGAGATGGTTTAACCGATGGT | TGGGAGCATCGGTTCTCTCGC | 2 |
| *DARK INDUCIBLE 11* (*DIN11*) | AT3G49620 | GTGGACGGTGATTGGATACC | TGGATTGGTAAACTCCGTTTG | 2 |
| *DON-GLUCOSYLTRANSFERASE 1* (*DOGT1*) | AT2G36800 | TGAGGGGATAACTGCTGGTC | CTGTTCAACCCCGGATCTTA | 2 |
| *EARLY LIGHT INDUCED PROTEIN 2* (*ELIP2*) | AT4G14690 | CACCACAAATGCCACAGTCT | TGCTAGTCTCCCGTTGATCC | 2 |
| *ELONGATED HYPOCOTYL 5* (*HY5*) | AT5G11260 | GAGGGAGGACACCGGCGGAG | CCTCTCTCTTGCTTGCTGAGCTGA | 2 |
| *PRODUCTION OF ANTHOCYANIN PIGMENT 1* (*PAP1*) | AT1G56650 | ACGCCCATTCCTACAACACCG | GGGCATTGAGATGGTTGCAGTCGT | 2 |
| *REPRESSOR OF UV-B PHOTOMORPHOGENESIS 2* (*RUP2*) | AT5G23730 | ATGGGCGCGTGATTCGCACG | ACGCGCCGTTTCTCCACACAG | 1.76 |
| *SOLANESYL DIPHOSPHATE SYNTHASE 1* (*SPS1*) | AT1G78510 | GCTCGGGAAGCCAGCAGGGA | TTGGCTCCCTCTCCAGAGCGA | 1.98 |
| *TYROSINE AMINOTRANSFERASE 3* (*TAT3*) | AT2G24850 | ATGATCGTCATGCCATCTCC | TGTGGACTTGTGGCATAGGA | 2 |
| Reference 1 | AT4G34270 | GTGAAAACTGTTGGAGAGAAGCAA | TCAACTGGATACCCTTTCGCA | 1.91 |
| Reference 2 | AT4G33380 | TTGAAAATTGGAGTACCGTACCAA | TCCCTCGTATACATCTGGCCA | 1.86 |
| Reference 3 | AT4G35510 | ACTTCCTCCGCGTCTCCATT | TTTATGTCCTGGCATTTCCAA | 1.97 |

Table S2. Summary of the ANOVA models used to estimate significant main effects of treatment, genotype, exposure time and the interactions (treatment × genotype, treatment × exposure time, genotype × exposure time, and treatment × genotype × exposure time) on transcript abundance measured by qRT-PCR in L*er*, *uvr8-2* and *cry1cry2* after 6 h and 12 h of treatments outdoors.

$df_{num}$ (numerator degrees of freedom), $df_{den}$ (denominator degrees of freedom).

| ***CHI* 6 h and 12 h** | | | | |
|---|---|---|---|---|
| **Source** | **$df_{num}$** | **$df_{den}$** | ***F*** | ***P*** |
| Treatment | 4 | 12 | 74.35 | <0.0001 |
| Genotype | 2 | 72 | 21.50 | <0.0001 |
| Exposure time | 1 | 72 | 2.39 | 0.1262 |
| Treatment × Genotype | 8 | 72 | 2.59 | 0.0152 |
| Treatment × Exposure time | 4 | 72 | 10.71 | <0.0001 |
| Genotype × Exposure time | 2 | 72 | 6.50 | 0.0025 |
| Treatment × Genotype × Exposure time | 8 | 72 | 2.15 | 0.0417 |
| ***CHI* 6 h** | | | | |
| Treatment | 4 | 12 | 62.06 | <0.0001 |
| Genotype | 2 | 29 | 7.92 | 0.0008 |
| Treatment × Genotype | 8 | 29 | 4.66 | 0.0001 |

| | | | | |
|---|---|---|---|---|
| **6 h L*er*** | | | | |
| Treatment | 4 | 12 | 55.50 | <0.0001 |
| **6 h *uvr8-2*** | | | | |
| Treatment | 4 | 12 | 23.64 | <0.0001 |
| **6 h *cry1cry2*** | | | | |
| Treatment | 4 | 12 | 45.99 | <0.0001 |
| ***CHI* 12 h** | | | | |
| Treatment | 4 | 12 | 27.89 | <0.0001 |
| Genotype | 2 | 27 | 38.48 | <0.0001 |
| Treatment × Genotype | 8 | 27 | 2.61 | 0.0295 |
| **12 h L*er*** | | | | |
| Treatment | 4 | 12 | 57.47 | <0.0001 |
| **12 h *uvr8-2*** | | | | |
| Treatment | 4 | 12 | 32.69 | <0.0001 |
| **12 h *cry1cry2*** | | | | |
| Treatment | 4 | 9 | 1.17 | 0.3844 |

| | | | | |
|---|---|---|---|---|
| ***CHS* 6 h and 12 h** | | | | |
| **Source** | | | | |
| Treatment | 4 | 12 | 140.11 | <0.0001 |
| Genotype | 2 | 70 | 41.78 | <0.0001 |
| Exposure time | 1 | 70 | 50.82 | <0.0001 |
| Treatment × Genotype | 8 | 70 | 27.83 | <0.0001 |
| Treatment × Exposure time | 4 | 70 | 41.49 | <0.0001 |
| Genotype × Exposure time | 2 | 70 | 11.21 | <0.0001 |
| Treatment × Genotype × Exposure time | 8 | 70 | 3.77 | 0.0010 |
| ***CHS* 6 h** | | | | |
| Treatment | 4 | 12 | 188.25 | <0.0001 |
| Genotype | 2 | 30 | 25.52 | <0.0001 |
| Treatment × Genotype | 8 | 30 | 21.38 | <0.0001 |
| **6 h L*er*** | | | | |
| Treatment | 4 | 12 | 228.11 | <0.0001 |
| **6 h *uvr8-2*** | | | | |
| Treatment | 4 | 12 | 44.53 | <0.0001 |
| **6 h *cry1cry2*** | | | | |
| Treatment | 4 | 12 | 139.22 | <0.0001 |
| ***CHS* 12 h** | | | | |
| Treatment | 4 | 12 | 45.86 | <0.0001 |
| Genotype | 2 | 25 | 26.52 | <0.0001 |
| Treatment × Genotype | 8 | 25 | 10.15 | <0.0001 |
| **12 h L*er*** | | | | |
| Treatment | 4 | 12 | 49.06 | <0.0001 |
| **12 h *uvr8-2*** | | | | |
| Treatment | 4 | 11 | 39.77 | <0.0001 |
| **12 h *cry1cry2*** | | | | |
| Treatment | 4 | 8 | 28.83 | <0.0001 |

| | | | | |
|---|---|---|---|---|
| ***DFR* 6 h and 12 h** | | | | |
| **Source** | | | | |
| Treatment | 4 | 12 | 25.78 | <0.0001 |
| Genotype | 2 | 72 | 42.95 | <0.0001 |
| Exposure time | 1 | 72 | 43.20 | <0.0001 |

| | | | | |
|---|---|---|---|---|
| Treatment × Genotype | 8 | 72 | 2.63 | 0.0137 |
| Treatment × Exposure time | 4 | 72 | 0.89 | 0.4729 |
| Genotype × Exposure time | 2 | 72 | 2.48 | 0.0912 |
| Treatment × Genotype × Exposure time | 8 | 72 | 0.94 | 0.4900 |
| ***DFR* 6 h** | | | | |
| Treatment | 4 | 12 | 20.36 | <0.0001 |
| Genotype | 2 | 30 | 41.28 | <0.0001 |
| Treatment × Genotype | 8 | 30 | 1.37 | 0.2500 |
| **6 h L*er*** | | | | |
| Treatment | 4 | 12 | 12.20 | 0.0003 |
| **6 h *uvr8-2*** | | | | |
| Treatment | 4 | 12 | 5.65 | 0.0086 |
| **6 h *cry1cry2*** | | | | |
| Treatment | 4 | 12 | 7.95 | 0.0023 |
| ***DFR* 12 h** | | | | |
| Treatment | 4 | 12 | 26.04 | <0.0001 |
| Genotype | 2 | 27 | 40.74 | <0.0001 |
| Treatment × Genotype | 8 | 27 | 4.49 | 0.0015 |
| **12 h L*er*** | | | | |
| Treatment | 4 | 12 | 31.26 | <0.0001 |
| **12 h *uvr8-2*** | | | | |
| Treatment | 4 | 12 | 10.57 | 0.0007 |
| **12 h *cry1cry2*** | | | | |
| Treatment | 4 | 9 | 9.64 | 0.0026 |

| | | | | |
|---|---|---|---|---|
| ***DIN11* 6 h and 12 h** | | | | |
| **Source** | | | | |
| Treatment | 4 | 12 | 1.88 | 0.1785 |
| Genotype | 2 | 71 | 18.63 | <0.0001 |
| Exposure time | 1 | 71 | 3.90 | 0.0521 |
| Treatment × Genotype | 8 | 71 | 1.78 | 0.0964 |
| Treatment × Exposure time | 4 | 71 | 3.06 | 0.0219 |
| Genotype × Exposure time | 2 | 71 | 1.65 | 0.1990 |
| Treatment × Genotype × Exposure time | 8 | 71 | 2.28 | 0.0315 |
| ***DIN11* 6 h** | | | | |
| Treatment | 4 | 12 | 3.86 | 0.0305 |
| Genotype | 2 | 28 | 14.74 | <0.0001 |
| Treatment × Genotype | 8 | 28 | 3.11 | 0.0122 |
| **6 h L*er*** | | | | |
| Treatment | 4 | 12 | 2.93 | 0.0663 |
| **6 h *uvr8-2*** | | | | |
| Treatment | 4 | 10 | 6.58 | 0.0073 |
| **6 h *cry1cry2*** | | | | |
| Treatment | 4 | 12 | 1.19 | 0.3640 |
| ***DIN11* 12 h** | | | | |
| Treatment | 4 | 12 | 0.55 | 0.7007 |
| Genotype | 2 | 28 | 11.90 | 0.0002 |
| Treatment × Genotype | 8 | 28 | 1.39 | 0.2437 |
| **12 h L*er*** | | | | |
| Treatment | 4 | 12 | 2.35 | 0.1133 |
| **12 h *uvr8-2*** | | | | |
| Treatment | 4 | 12 | 1.58 | 0.2427 |

| | | | | |
|---|---|---|---|---|
| **12 h *cry1cry2*** | | | | |
| Treatment | 4 | 10 | 0.72 | 0.598 |

| | | | | |
|---|---|---|---|---|
| ***DOGT1* 6 h and 12 h** | | | | |
| **Source** | | | | |
| Treatment | 4 | 12 | 39.62 | <0.0001 |
| Genotype | 2 | 70 | 26.77 | <0.0001 |
| Exposure time | 1 | 70 | 0.03 | 0.8552 |
| Treatment × Genotype | 8 | 70 | 2.52 | 0.0180 |
| Treatment × Exposure time | 4 | 70 | 3.60 | 0.0100 |
| Genotype × Exposure time | 2 | 70 | 0.58 | 0.5616 |
| Treatment × Genotype × Exposure time | 8 | 70 | 2.05 | 0.0525 |
| ***DOGT1* 6 h** | | | | |
| Treatment | 4 | 12 | 34.51 | <0.0001 |
| Genotype | 2 | 27 | 16.90 | <0.0001 |
| Treatment × Genotype | 8 | 27 | 1.94 | 0.0954 |
| **6 h L*er*** | | | | |
| Treatment | 4 | 12 | 31.25 | <0.0001 |
| **6 h *uvr8-2*** | | | | |
| Treatment | 4 | 9 | 6.61 | 0.0091 |
| **6 h *cry1cry2*** | | | | |
| Treatment | 4 | 12 | 7.34 | 0.0031 |
| ***DOGT1* 12 h** | | | | |
| Treatment | 4 | 12 | 12.72 | 0.0003 |
| Genotype | 2 | 28 | 14.57 | <0.0001 |
| Treatment × Genotype | 8 | 28 | 3.18 | 0.0107 |
| **12 h L*er*** | | | | |
| Treatment | 4 | 12 | 8.18 | 0.0020 |
| **12 h *uvr8-2*** | | | | |
| Treatment | 4 | 12 | 5.94 | 0.0071 |
| **12 h *cry1cry2*** | | | | |
| Treatment | 4 | 10 | 5.39 | 0.0141 |

| | | | | |
|---|---|---|---|---|
| ***ELIP2* 6 h and 12 h** | | | | |
| **Source** | | | | |
| Treatment | 4 | 12 | 234.74 | <0.0001 |
| Genotype | 2 | 66 | 48.40 | <0.0001 |
| Exposure time | 1 | 66 | 161.90 | <0.0001 |
| Treatment × Genotype | 8 | 66 | 21.61 | <0.0001 |
| Treatment × Exposure time | 4 | 66 | 7.93 | <0.0001 |
| Genotype × Exposure time | 2 | 66 | 3.81 | 0.0272 |
| Treatment × Genotype × Exposure time | 8 | 66 | 1.88 | 0.0772 |
| ***ELIP2* 6 h** | | | | |
| Treatment | 4 | 11 | 150.34 | <0.0001 |
| Genotype | 2 | 26 | 15.01 | <0.0001 |
| Treatment × Genotype | 8 | 26 | 13.94 | <0.0001 |
| **6 h L*er*** | | | | |
| Treatment | 4 | 10 | 100.71 | <0.0001 |
| **6 h *uvr8-2*** | | | | |
| Treatment | 4 | 10 | 72.81 | <0.0001 |
| **6 h *cry1cry2*** | | | | |
| Treatment | 4 | 11 | 46.77 | <0.0001 |

| | | | | |
|---|---|---|---|---|
| ***ELIP2* 12 h** | | | | |
| Treatment | 4 | 12 | 153.42 | <0.0001 |
| Genotype | 2 | 27 | 37.17 | <0.0001 |
| Treatment × Genotype | 8 | 27 | 10.21 | <0.0001 |
| **12 h L*er*** | | | | |
| Treatment | 4 | 12 | 146.00 | <0.0001 |
| **12 h *uvr8-2*** | | | | |
| Treatment | 4 | 12 | 213.36 | <0.0001 |
| **12 h *cry1cry2*** | | | | |
| Treatment | 4 | 9 | 41.08 | <0.0001 |

| | | | | |
|---|---|---|---|---|
| ***HY5* 6 h and 12 h** | | | | |
| **Source** | | | | |
| Treatment | 4 | 12 | 4.24 | 0.0228 |
| Genotype | 2 | 71 | 1.64 | 0.2017 |
| Exposure time | 1 | 71 | 5.50 | 0.0218 |
| Treatment × Genotype | 8 | 71 | 1.03 | 0.4203 |
| Treatment × Exposure time | 4 | 71 | 0.21 | 0.9294 |
| Genotype × Exposure time | 2 | 71 | 0.15 | 0.8600 |
| Treatment × Genotype × Exposure time | 8 | 71 | 0.81 | 0.5987 |
| ***HY5* 6 h** | | | | |
| Treatment | 4 | 12 | 48.74 | <0.0001 |
| Genotype | 2 | 29 | 9.78 | 0.0006 |
| Treatment × Genotype | 8 | 29 | 3.51 | 0.0059 |
| **6 h L*er*** | | | | |
| Treatment | 4 | 12 | 28.83 | <0.0001 |
| **6 h *uvr8-2*** | | | | |
| Treatment | 4 | 11 | 16.25 | 0.0001 |
| **6 h *cry1cry2*** | | | | |
| Treatment | 4 | 12 | 8.03 | 0.0022 |
| ***HY5* 12 h** | | | | |
| Treatment | 4 | 12 | 1.83 | 0.1880 |
| Genotype | 2 | 27 | 1.81 | 0.1836 |
| Treatment × Genotype | 8 | 27 | 1.59 | 0.1742 |
| **12 h L*er*** | | | | |
| Treatment | 4 | 12 | 1.69 | 0.2156 |
| **12 h *uvr8-2*** | | | | |
| Treatment | 4 | 11 | 1.74 | 0.2111 |
| **12 h *cry1cry2*** | | | | |
| Treatment | 4 | 10 | 4.19 | 0.0302 |

| | | | | |
|---|---|---|---|---|
| ***PAP1* 6 h and 12 h** | | | | |
| **Source** | | | | |
| Treatment | 4 | 12 | 12.54 | 0.0003 |
| Genotype | 2 | 70 | 103.72 | <0.0001 |
| Exposure time | 1 | 70 | 4.10 | 0.0468 |
| Treatment × Genotype | 8 | 70 | 1.61 | 0.1391 |
| Treatment × Exposure time | 4 | 70 | 3.11 | 0.0206 |
| Genotype × Exposure time | 2 | 70 | 4.81 | 0.0111 |
| Treatment × Genotype × Exposure time | 8 | 70 | 1.51 | 0.1695 |
| ***PAP1* 6 h** | | | | |

| | | | | |
|---|---|---|---|---|
| Treatment | 4 | 12 | 10.76 | 0.0006 |
| Genotype | 2 | 29 | 84.15 | <0.0001 |
| Treatment × Genotype | 8 | 29 | 3.20 | 0.0100 |
| **6 h L*er*** | | | | |
| Treatment | 4 | 12 | 10.96 | 0.0006 |
| **6 h *uvr8-2*** | | | | |
| Treatment | 4 | 11 | 3.83 | 0.0346 |
| **6 h *cry1cry2*** | | | | |
| Treatment | 4 | 12 | 19.68 | <0.0001 |
| ***PAP1* 12 h** | | | | |
| Treatment | 4 | 12 | 6.60 | 0.0048 |
| Genotype | 2 | 26 | 61.37 | <0.0001 |
| Treatment × Genotype | 8 | 26 | 1.32 | 0.2761 |
| **12 h L*er*** | | | | |
| Treatment | 4 | 12 | 30.62 | <0.0001 |
| **12 h *uvr8-2*** | | | | |
| Treatment | 4 | 11 | 5.14 | 0.0139 |
| **12 h *cry1cry2*** | | | | |
| Treatment | 4 | 9 | 1.55 | 0.2680 |

| ***RUP2* 6 h and 12 h** | | | | |
|---|---|---|---|---|
| **Source** | | | | |
| Treatment | 4 | 12 | 119.28 | <0.0001 |
| Genotype | 2 | 72 | 3.48 | 0.0361 |
| Exposure time | 1 | 72 | 5.30 | 0.0242 |
| Treatment × Genotype | 8 | 72 | 5.54 | <0.0001 |
| Treatment × Exposure time | 4 | 72 | 10.90 | <0.0001 |
| Genotype × Exposure time | 2 | 72 | 6.26 | 0.0031 |
| Treatment × Genotype × Exposure time | 8 | 72 | 3.16 | 0.0040 |
| ***RUP2* 6 h** | | | | |
| Treatment | 4 | 12 | 101.16 | <0.0001 |
| Genotype | 2 | 30 | 2.65 | 0.0868 |
| Treatment × Genotype | 8 | 30 | 8.84 | <0.0001 |
| **6 h L*er*** | | | | |
| Treatment | 4 | 12 | 120.54 | <0.0001 |
| **6 h *uvr8-2*** | | | | |
| Treatment | 4 | 12 | 43.77 | <0.0001 |
| **6 h *cry1cry2*** | | | | |
| Treatment | 4 | 12 | 33.86 | <0.0001 |
| ***RUP2* 12 h** | | | | |
| Treatment | 4 | 12 | 53.16 | <0.0001 |
| Genotype | 2 | 27 | 9.56 | 0.0007 |
| Treatment × Genotype | 8 | 27 | 3.33 | 0.0088 |
| **12 h L*er*** | | | | |
| Treatment | 4 | 11 | 52.65 | <0.0001 |
| **12 h *uvr8-2*** | | | | |
| Treatment | 4 | 12 | 39.48 | <0.0001 |
| **12 h *cry1cry2*** | | | | |
| Treatment | 4 | 10 | 2.54 | 0.1053 |

| ***SPS1* 6 h and 12 h** |
|---|
| **Source** |

| | | | | |
|---|---|---|---|---|
| Treatment | 4 | 12 | 84.76 | <0.0001 |
| Genotype | 2 | 72 | 2.04 | 0.1374 |
| Exposure time | 1 | 72 | 1.80 | 0.1844 |
| Treatment × Genotype | 8 | 72 | 4.02 | 0.0005 |
| Treatment × Exposure time | 4 | 72 | 14.32 | <0.0001 |
| Genotype × Exposure time | 2 | 72 | 7.35 | 0.0012 |
| Treatment × Genotype × Exposure time | 8 | 72 | 5.30 | <0.0001 |
| ***SPS1* 6 h** | | | | |
| Treatment | 4 | 12 | 78.42 | <0.0001 |
| Genotype | 2 | 30 | 3.36 | 0.0483 |
| Treatment × Genotype | 8 | 30 | 7.39 | <0.0001 |
| **6 h L*er*** | | | | |
| Treatment | 4 | 12 | 57.53 | <0.0001 |
| **6 h *uvr8-2*** | | | | |
| Treatment | 4 | 12 | 49.54 | <0.0001 |
| **6 h *cry1cry2*** | | | | |
| Treatment | 4 | 12 | 34.96 | <0.0001 |
| ***SPS1* 12 h** | | | | |
| Treatment | 4 | 12 | 24.81 | <0.0001 |
| Genotype | 2 | 27 | 8.47 | 0.0014 |
| Treatment × Genotype | 8 | 27 | 6.67 | 0.0001 |
| **12 h L*er*** | | | | |
| Treatment | 4 | 11 | 37.21 | <0.0001 |
| **12 h *uvr8-2*** | | | | |
| Treatment | 4 | 12 | 26.34 | <0.0001 |
| **12 h *cry1cry2*** | | | | |
| Treatment | 4 | 10 | 2.70 | 0.0925 |

| | | | | |
|---|---|---|---|---|
| ***TAT3* 6 h and 12 h** | | | | |
| **Source** | | | | |
| Treatment | 4 | 12 | 4.01 | 0.0271 |
| Genotype | 2 | 70 | 4.67 | 0.0125 |
| Exposure time | 1 | 70 | 0.25 | 0.6220 |
| Treatment × Genotype | 8 | 70 | 1.67 | 0.1210 |
| Treatment × Exposure time | 4 | 70 | 0.66 | 0.6238 |
| Genotype × Exposure time | 2 | 70 | 0.30 | 0.7433 |
| Treatment × Genotype × Exposure time | 8 | 70 | 1.12 | 0.3637 |
| ***TAT3* 6 h** | | | | |
| Treatment | 4 | 12 | 4.31 | 0.0218 |
| Genotype | 2 | 28 | 5.85 | 0.0075 |
| Treatment × Genotype | 8 | 28 | 1.93 | 0.0942 |
| **6 h L*er*** | | | | |
| Treatment | 4 | 12 | 1.14 | 0.3857 |
| **6 h *uvr8-2*** | | | | |
| Treatment | 4 | 11 | 3.74 | 0.0371 |
| **6 h *cry1cry2*** | | | | |
| Treatment | 4 | 11 | 4.53 | 0.0208 |
| ***TAT3* 12 h** | | | | |
| Treatment | 4 | 12 | 2.43 | 0.1046 |
| Genotype | 2 | 27 | 2.37 | 0.1123 |
| Treatment × Genotype | 8 | 27 | 1.71 | 0.1424 |
| **12 h L*er*** | | | | |

| | | | | |
|---|---|---|---|---|
| Treatment | 4 | 11 | 1.55 | 0.2557 |
| **12 h *uvr8-2*** | | | | |
| Treatment | 4 | 11 | 1.55 | 0.2552 |
| **12 h *cry1cry2*** | | | | |
| Treatment | 4 | 11 | 2.05 | 0.1560 |

**Methods S1**

***Description of the filters and the waveband contrasts***

The treatments made use of five long-pass filters differing in cut-off wavelength (Figure 1). UV wavelengths $\lambda > 290$ nm were all transmitted by clear acrylic PLEXIGLAS 2458 GT (Evonik, Essen, Germany), $\lambda < 315$ nm were excluded with PLEXIGLAS 2458 GT (Evonik) overlaid with polyester film (Autostat CT-5 MacDermid Autotype, Wantage, UK), $\lambda < 350$ nm were excluded by solar Clear acrylic PLEXIGLAS 0Z023 GT (Evonik), $\lambda < 400$ nm were excluded by clear polycarbonate Makrolife (Arla Plast, Borensberg, Sweden) and $\lambda < 500$ nm were excluded by yellow acrylic PLEXIGLAS 1C33 GT (Evonik) (Figure 1). The effect of UV-B radiation was tested from contrasts between the treatments >290 nm versus >315 nm, the effect of UV-A radiation from contrasts between >315 nm versus >400 nm, the effect of UV-A short wavelength radiation (UV-$A_{sw}$) from contrasts between >315 nm versus >350 nm, the effect of UV-A long wavelength radiation (UV-$A_{lw}$) from contrasts between >350 nm versus >400 nm, and the effect of blue light from contrasts between >400 nm versus >500 nm. In addition, we also tested the effect of wavelength $\lambda < 350$ nm from contrasts >290 nm versus >350 nm and the effect of wavelength $\lambda > 350$ nm from contrasts >350 nm versus >500 nm (Figure 1).
